# Brain-wide mapping reveals temporal and sexually dimorphic opioid actions

**DOI:** 10.1101/2025.02.19.638902

**Authors:** Iaroslavna Vasylieva, Megan C. Smith, Eshan Aravind, Kelin He, Tianhan Ling, Jenesis Kozel, Stephanie Puig, Katarzyna M. Kedziora, Jessica J. Scarlett, Paul N. Joseph, Matthew D. Lycas, Ulrik Gether, Ryan W. Logan, Zachary Freyberg, Alan M. Watson

## Abstract

While the molecular and cellular effects of opioids have been extensively studied, the precise mechanisms by which these drugs target specific brain regions over time remain unclear. Similarly, despite well-documented sex differences in opioid responses, the anatomical basis for this sexual dimorphism is not well characterized. To address these questions, we developed an automated, scalable, and unbiased approach for whole-brain anatomical mapping of the neuronal activity marker c-Fos in response to acute morphine exposure. Using ribbon scanning confocal microscopy, we imaged whole cleared brains from male and female wild-type mice at 1 hour and 4 hours post-morphine administration. Our whole-brain analysis of c-Fos expression revealed distinct patterns of morphine-induced regional brain activation across time and sex. Notably, we observed a greater number of structures with significant activity differences at 4 hours compared to 1 hour. In male mice, significant changes were primarily localized to regions within the dopamine system, whereas in female mice, they were concentrated in cortical regions. By combining high-throughput imaging with whole-brain expression analysis, particularly in the context of opioid actions, our approach provides a more comprehensive understanding of how drugs of abuse affect the brain.

## Introduction

Opioids are some of the most used and abused substances today, playing a central role in the treatment of pain. However, these drugs can also induce tolerance and dependence, contributing significantly to their high potential for abuse^1^. As a result, opioid abuse prevalence has skyrocketed in the United States and globally^2–4^. Current pharmacological treatments for opioid use disorder (OUD) are effective for mitigating cravings, although ∼90% of patients relapse within several months of treatment cessation^5^. Thus, to identify novel treatments for OUD and interventions capable of slowing the opioid epidemic, we must elucidate the neurobiology of opioid addiction and its relationships to factors that contribute to craving and relapse vulnerability Initial, acute exposure to opioids may lead to rapid adaptive changes in the brain that reflect the early mechanisms of opioid dependence and risk of subsequent addiction. Indeed, treating humans or rodents with opioid antagonists, such as naltrexone, elicits signs of withdrawal from opioids^6^. Changes in neuronal activity across various populations of neurons occur following acute administration of opioids, likely underlying the adaptive changes in the brain that contribute to tolerance and withdrawal^7–10^. Revealing where these changes in activity occur and their relationship to drug-related behaviors is critical for understanding the fundamental actions of drugs of abuse, including opioids, and for providing potential neurobiological targets for therapeutic intervention. Consequently, the neuroscience field has been grappling with how to effectively investigate activity across multiple scales within the same brain, from the single cell to ensembles to entire brain regions, and *how* drugs modulate these changes in activity.

While acknowledging the biological importance of neuronal heterogeneity to addiction and the need for scalability, most current approaches are still limited by their ability to resolve distinct changes in activity in response to drugs of abuse at the single-cell level within larger populations across the whole brain. For example, immunolabeling of neuronal activity markers across various brain regions has been valuable for revealing new neurobiological substrates of complex drug-related behaviors^7,9,11,12^. Yet, because of various constraints of throughput and resolution, earlier work has typically focused on investigating discrete regions in areas previously implicated in addiction. As a result, these prior limitations have created a bottleneck for the development of a more integrated understanding of the biological underpinnings of addiction at a whole brain level. The ability to widely survey the whole brain at sufficient resolution permits us to move beyond single brain regions. Rather, we can commence examining current opioid actions across multiple regions – a key next step in the generation of a holistic, more integrated understanding of drug actions.

We have developed a workflow to address these challenges which offer the capability to readily move across scales throughout whole intact brains in three dimensions (3D). Rather than being limited to a single brain region or smaller subsets of neurons, newer whole brain imaging and analysis methods provide the opportunity to probe drug-induced effects in an unbiased, discovery-based manner. Herein we used tissue clearing^13^ and high-speed ribbon scanning confocal microscopy^14^ (RSCM) to collect highresolution imagery of whole brains and applied a novel automated high-performance computing workflow that detects and maps cells within the brain. The development of this integrated workflow allowed us to digitally reconstruct entire adult mouse brains at sub-cellular resolution and therefore to comprehensively map populations of activated neurons across the entire brain in dozens of animals. This automated approach to data analysis allows us to tackle experimental questions like time-, sex-, and brain region-dependent differences when examining opioid actions.

We utilized our workflow to comprehensively characterize the global anatomical expression patterns of the immediate early gene (IEG) *c-Fos*, a classic neuronal activity marker, in response to the administration of the prototypical opioid, morphine. Our data demonstrated broad patterns of increased activity within the brains of morphine-exposed wildtype mice. The patterns of morphine-induced activity demonstrated distinct differences in neuronal activity across time of morphine exposure as well as across male and female animals. Specifically, we find that the mesolimbic dopamine system is activated more strongly in male mice, whereas cortical regions are activated more strongly in female mice. In summary, our imaging and analysis approaches open the avenue for answering long-standing questions in opioid actions, including resolving cellular heterogeneity within larger populations and mapping their changes in response to drug exposure. Following peer review, at the time of publication, and in line with FAIR^15,16^ principles, we will make our tools openly available by depositing a comprehensive set of code, microscopy data and examples into public repositories so that the community can reproduce our work or apply our methods to novel questions.

## Results

### Development of a whole brain imaging and analysis workflow

To investigate the impact of different durations of opioid exposure on the activities of cell subpopulations throughout the whole adult mouse brain, we administered morphine to male and female wild-type mice versus a vehicle control (Fig. 1a). 1h or 4h following exposure, brains were collected and subsequently fluorescently stained for c-Fos, followed by optical clearing. We then imaged the cleared and c-Fos-labeled brains via RSCM. After imaging, data from each brain was processed through an automated analysis pipeline (Fig. 1b) which consisted of parallel steps for down-scaling the data for alignment to the Allen Mouse Brain Common Coordinate Framework version 3 (CCFv3^17^) and detection of c-Fos-positive cells throughout the brain. We used a combination of conventional and deep-learning spot detection approaches followed by clustering duplicate detections and neural network-based discrimination of true cells and artifacts.

**Figure 1.**
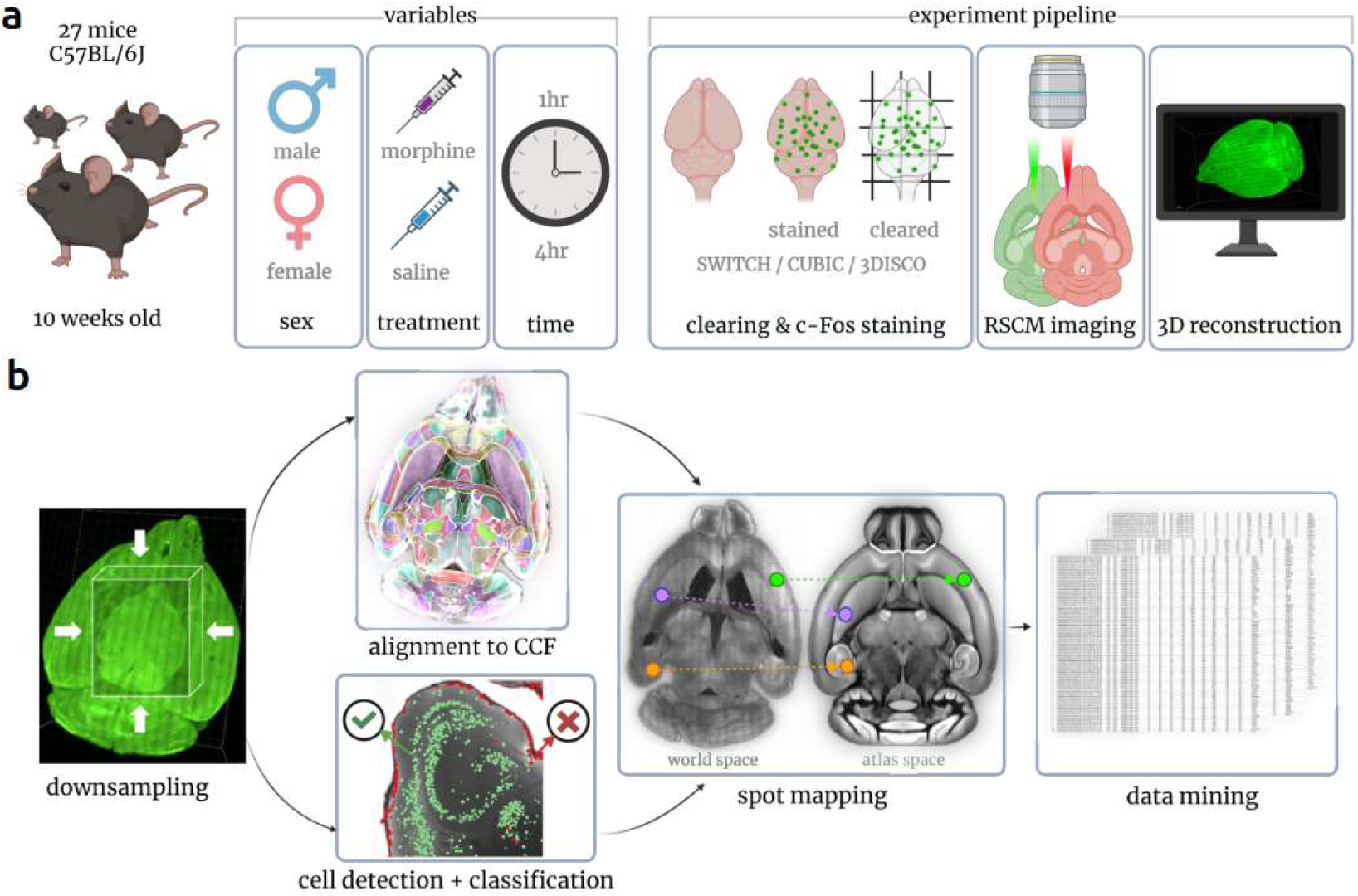
Experimental design. **(a)** 10-week-old male and female mice were injected with morphine or saline. 1h or 4h post injection, their brains were harvested, stained for c-Fos and rendered optically transparent. Cleared brains were imaged using RSCM then reconstructed in 3D. The 3D images were run through the quantitative data analysis pipeline **(b)**. The 3D brain reconstructions were first downsampled to ∼10 μm resolution. Images were subsequently registered to the CCFv3. In parallel, c-Fos-positive cells were detected and each location recorded in the 3D dataset (world space). Each spot was then classified as cell / non-cell using a neural network. Coordinates for each cell were transformed to the CCFv3 space using the deformation field obtained from the image registration procedure and associated with a CCFv3 brain region (spot mapping). The resulting tables of spot centroids and corresponding brain regions were aggregated by brain region and across treatment groups for statistical analysis.

All aspects of the computational workflow were completed semi-autonomously on local high-performance computing resources managed by the Simple Linux Utility for Resource Management (SLURM^18^). Following RSCM, data was automatically queued for stitching and assembly into 3D datasets. After manual verification of each 3D dataset, spot detection, alignment and classification were queued as a single job, outputting intermediate processing products like the alignment deformation field and finally the spot tables which included CCFv3 mapped structures. Where possible, we reused already available open-source toolkits, developing novel means to knit the software together and produce an automated workflow requiring minimal intervention. Following an initial pass, the results of each alignment and cell classification were manually reviewed and when appropriate, reprocessed with modified parameters. Although we used SLURM to manage this process, the workflow can be run on a single mid-to high-end workstation with a NVIDIA GPU making it accessible to most research labs.

The performance of automated spot detection and classification was measured against a ground truth data sample collected for each brain. Acceptance criterion was f1-score >80%. For the brains having a lower f1-score, the base classification model was fine-tuned to reach the 80% threshold. For each brain, the coordinates of c-Fos-positive cells were transformed into CCFv3 space and exported as CSV files that would be the basis for all subsequent data mining. During analysis, we took an unbiased approach by inspecting morphine’s effects on every brain structure at every depth level in the CCFv3 structure tree. This included calculating the total numbers and densities of c-Fospositive cells for each respective brain structure. We also calculated mean cell densities for each structure, for “all morphine” and “all saline” groups separately, and then calculated the morphine/saline ratio along with corresponding p-values. We hereafter refer to the density ratio simply as density fold-change.

### Morphine induces broad increases in c-Fos activity across the whole brain

To determine whether morphine exposure induced measurable changes in c-Fos expression across the mouse brain, we plotted coronal heatmaps of the average cellular density of c-Fos-positive cells from saline-(Fig. 2a) and morphine-exposed (Fig. 2b) animals at 1mm projections. There were clearly observable increases in c-Fos-positive cell density across the brains of morphine-exposed animals with particularly apparent increases in the cortical layers. Quantitative analysis confirmed that more c-Fos-positive cells were present in brains of morphine-exposed mice compared to brains of saline-treated controls. This increase in morphine-induced activity was observed across most brain regions (Fig. 2c) and remained consistent overall when examining individual experimental groups (Fig. 3). Out of 840 structures in the CCFv3, there were 713 structures with an increase in c-Fos-positive cells following morphine exposure. In contrast, 125 structures showed a decrease in c-Fos-positive cells, and 2 structures showed no change following either treatment. Of the structures with decreased activity, the highest numbers were in the Isocortex (52 structures), Thalamus (17 structures) and Midbrain (12 structures). In the Thalamus, ∼25% of substructures showed reduced activity.

**Figure 2.**
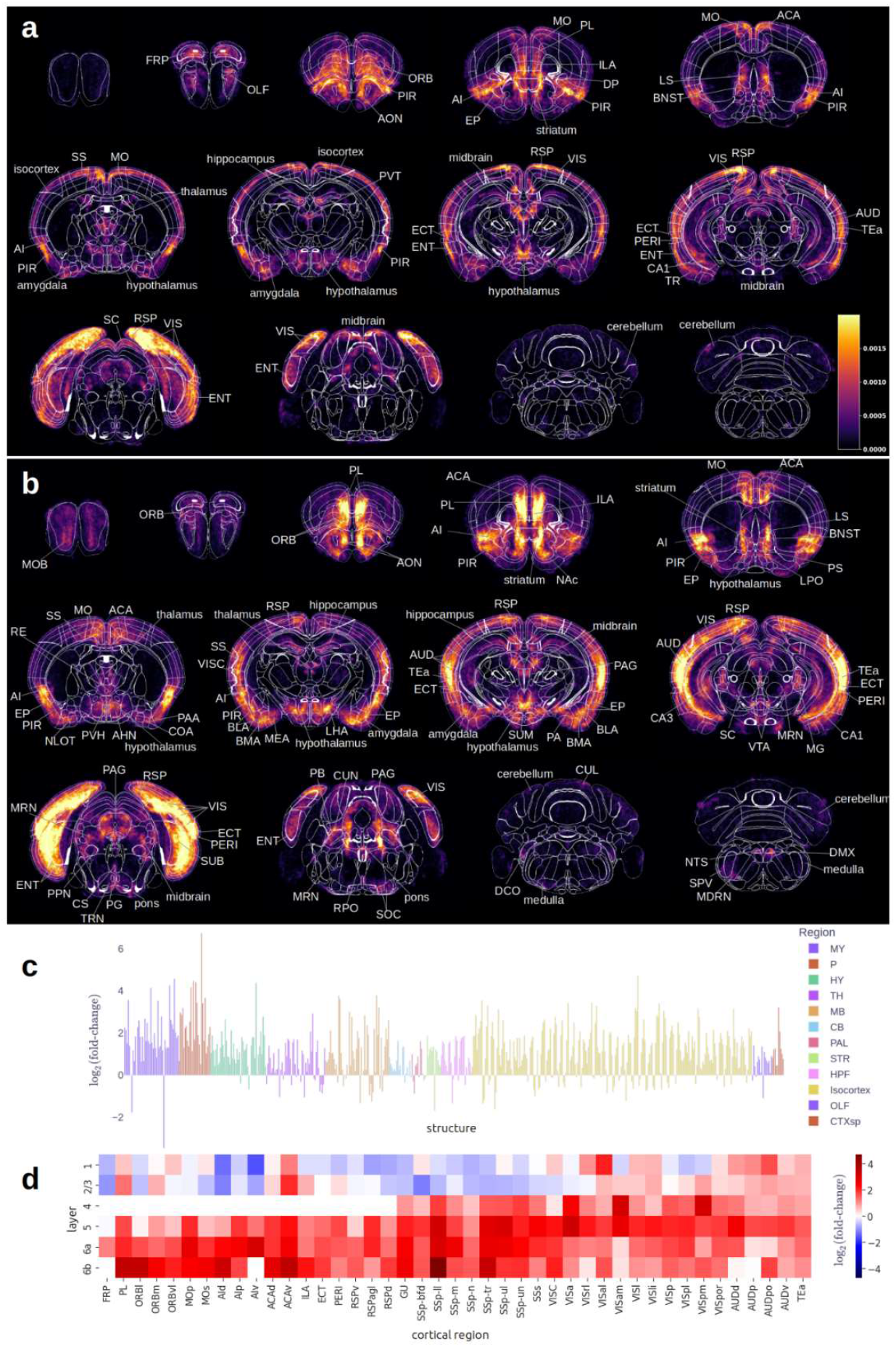
Global c-Fos expression in mice exposed to morphine. 1mm thick coronal heatmaps^*^ of c-Fos positive neurons for all **a)** saline and **b)** morphine treated animals. Areas containing the brightest signal are labeled according to the CCFv3 structure. Morphine/saline c-Fos density fold-change is displayed for all grey matter brain structures **c)** within the CCFv3 and grouped by larger brain region as indicated by color. Cortical areas **d)** are grouped by layer with color representing log2(fold-change). Blue indicates depressed activity in a region due to morphine exposure and red indicates enhanced activity. ^*^Heatmaps in panels a and b were calculated by aggregating all c-FOS positive neurons, dividing by the total number of animals and then smoothing with a gaussian kernel. All images are displayed on the same scale (CCFv3).

**Figure 3.**
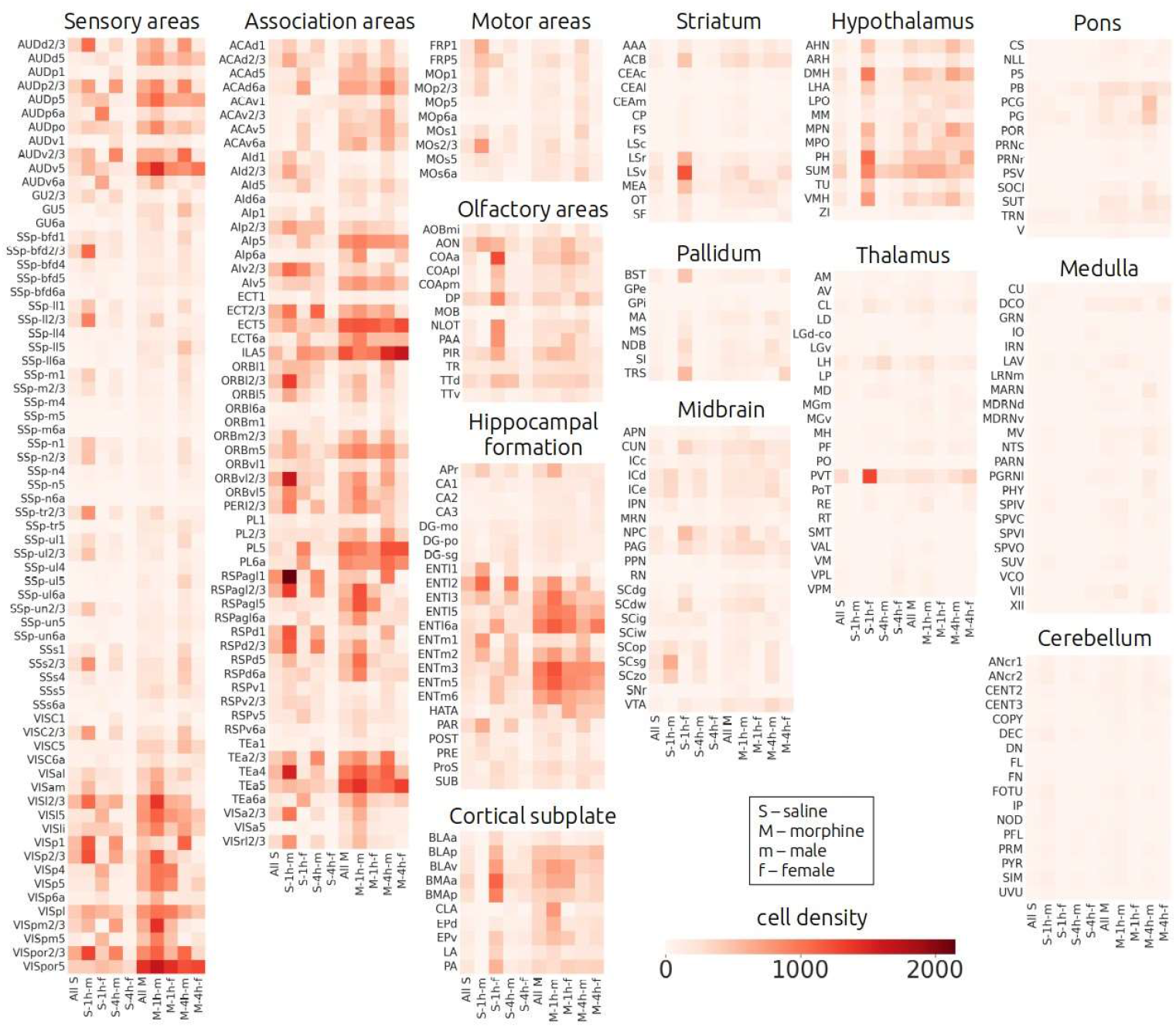
c-Fos-positive cell densities. A heat map displaying the density of c-Fos expressing cells (cells/mm^3^) organized according to experimental groups and arranged according to prominent brain structures. Structures were selected based on their depth in the CCFv3 structure tree (leaves, deepest structures) and volume > 0.25 mm^3^. Experimental groups were the smallest sets of animals that differed by all 3 variables (treatment, time-point, sex). There were 8 experimental groups: morphine-1h-male (M-1h-m, N=5), morphine-1h-female (M-1h-f, N=3), morphine-4h-male (M-4h-m, N=4), morphine-4h-female (M-4h-f, N=3), saline-1h-male (S-1h-m, N=3), saline-1h-female (S-1h-f, N=2), saline-4h-male (S-4h-m, N=3), saline-4h-female (S-4h-f, N=4)

At a coarser scale, we examined large brain regions that collectively constitute 90% of the brain volume. Our data showed that each of these structures on average had more c-Fos-positive cells in the brain from morphine-exposed animals compared to saline controls (Table 1, Fig. S1). Morphine-treated brains demonstrated the greatest fold-change in Pons and Medulla, and the lowest fold-change in the Thalamus and Cerebellum. Statistically significant increases at this depth of the structure tree were only observed in the Pons, Cortical Subplate and Hippocampal Formation. Overall, the highest density of morphine-activated c-Fos puncta was observed in the Cortical subplate and Isocortex. Most of the large structures contained subregions with both decreased and increased c-Fos signal. For example, the Isocortex showed mostly increased signal in layers 4-6, and decreased signal in cortical layers 1-3 (Fig. 2d). However, layers 1-3 displayed increased c-Fos signal in regions associated with auditory and visual processing.

**Table 1.** c-Fos cell densities on major brain structures. *P<0.05

| STRUCTURE | DENSITY MORPHINE<br>[CELLS/MM<sup>3</sup>] | DENSITY SALINE<br>[CELLS/MM<sup>3</sup>] | DENSITY<br>FOLD CHANGE | P-VALUE |
| --- | --- | --- | --- | --- |
| PONS | 121.47 | 32.28 | 3.76 | 0.0297 * |
| MEDULLA | 48.93 | 17.46 | 2.80 | 0.1524 |
| CORTICAL SUBPLATE | 463.94 | 212.38 | 2.18 | 0.0301 * |
| HYPOTHALAMUS | 331.58 | 163.64 | 2.03 | 0.1069 |
| HIPPOCAMPAL<br>FORMATION | 393.18 | 215.53 | 1.82 | 0.0376 * |
| STRIATUM | 119.69 | 71.3 | 1.68 | 0.2260 |
| PALLIDUM | 121.84 | 74.6 | 1.63 | 0.2921 |
| ISOCORTEX | 417.89 | 256.76 | 1.63 | 0.2879 |
| MIDBRAIN | 180.87 | 119.48 | 1.51 | 0.1869 |
| OLFACTORY AREAS | 329.66 | 222.46 | 1.48 | 0.2126 |
| CEREBELLUM | 49.68 | 39.4 | 1.26 | 0.6441 |
| THALAMUS | 63.82 | 52.47 | 1.22 | 0.6373 |

Of the smaller structures within the CCFv3, 46 exhibited significant differences after morphine administration (p<0.05) and met other relevant criteria including: 1) gray matter composition, 2) >0.25 mm^3^ volume (to reduce error imposed from registration artifacts), and 3) possessed ≥25 c-Fos spots detected on average in brains of both morphine and saline groups (Fig. 4a, Table S1). Among these structures approximately half (24) were within the Isocortex including the Anterior cingulate area (ACA), Auditory areas (AUD), Visual areas (VIS), Somatosensory areas (SS), Prelimbic area (PL), Temporal association area (TEa), Visceral area (VISC), Ectorhinal area (ECT), Infralimbic area (ILA), and the Agranular Insular Area (AI). The remaining structures were found within the Hippocampal Formation, Midbrain including the Ventral Tegmental Area (VTA), Cortical Subplate, Pons and Cerebellum. In addition, there were 31 sufficiently large structures (>0.25 mm^3^, >25 cells) that approached significance (0.05 <= p < 0.1, Table S2). These included the Nucleus Accumbens (ACB, p=0.07), periaqueductal grey (PAG^19^, p=0.07) and the midbrain Raphe nuclei (RAmb, p=0.06) which have been previously shown^20^ to respond following morphine exposure^21^. Due to the hierarchical nature of the CCFv3, a child structure having a statistically significant difference would often cause its parent structures to have a statistically significant difference. For simplicity, we excluded such parent structures from the above lists (see Table S8 for a full list of structures and their significance).

**Figure 4.**
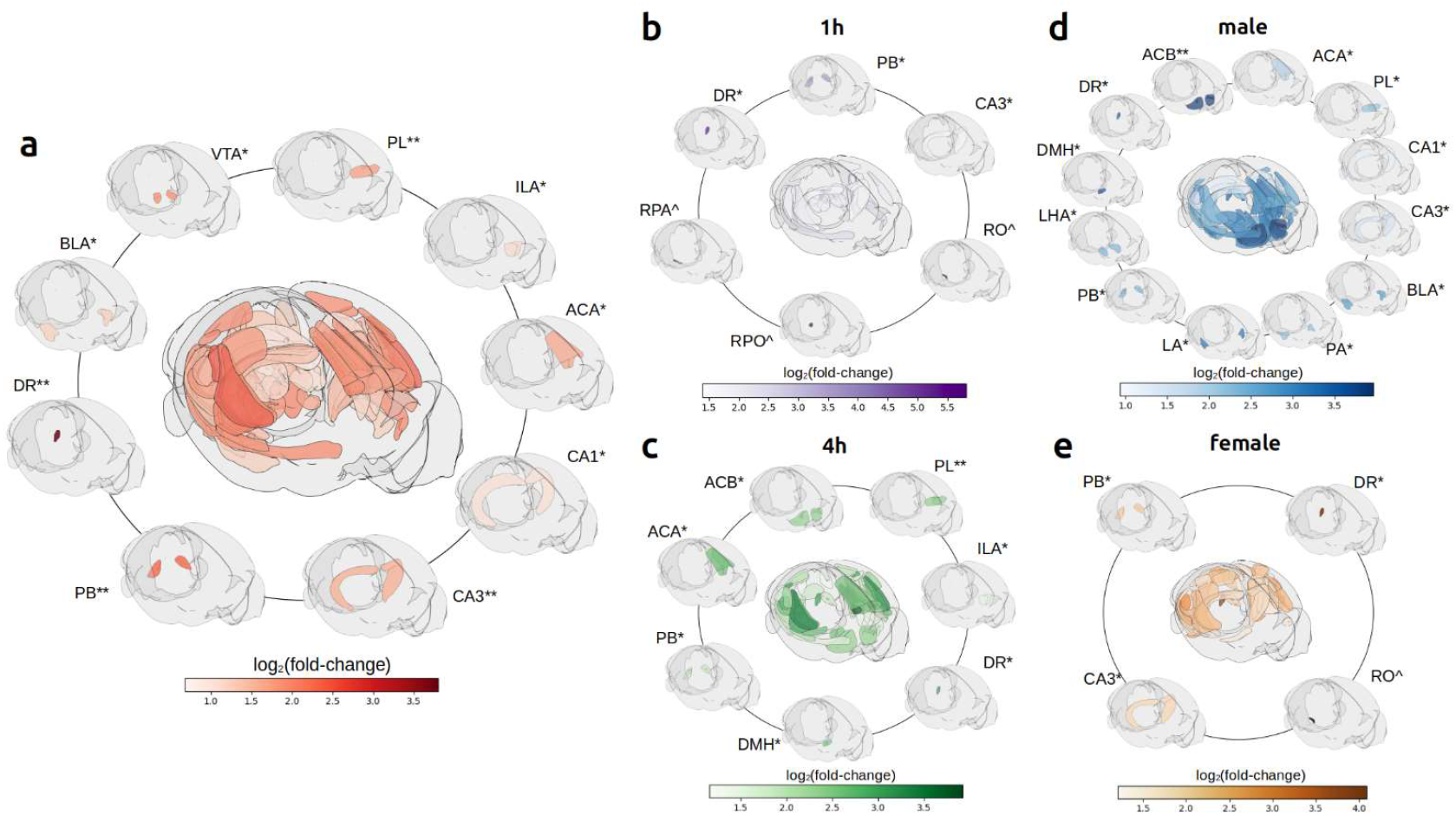
Structures with significant differences in c-Fos expression following morphine exposure. Displayed are brain regions that show significant differences between saline control and morphine exposed animals. Structures mentioned in the literature as implicated in opioid addiction are indicated along the outer circle whereas all significant structures are represented in the middle. Panel a) represents 46 structures (p<0.05) between all morphine and all saline animals from Table S1. Time points and sex differences are compared in panels b-c (tables S4,S5) and d-e (tables S6, S7), respectively. Color represents logarithm of density fold-change. * - p < 0.05, ** - p <0.01, -^ binary effect. Structures with binary effects are shown in black. Dorsal Raphe nucleus falls below the size threshold, however it is included because it displays a statistically significant difference in all subgroups (see Table S8). Other Raphe nuclei are shown since in many cases they had binary effects (RPO, RPA, RO).

We observed that many small structures only contained c-Fos-positive puncta in either the morphine exposed animals or the saline controls (density=0). We hereafter refer to these structures as having a binary effect (Table S3, Fig. S3a). When calculating fold-change, the binary structures were excluded due to the undefined nature of dividing by 0. Most of these structures fell below our size threshold but were intriguing because in most of the structures, c-Fos activity was observed exclusively in response to morphine treatment. Among all morphine and all saline brains, morphine brains exhibited binary effect in a total of 39/41 structures: in the Medulla, Isocortex, Midbrain, Pons, Hypothalamus and Thalamus (ordered by decreasing number of “binary” structures). In contrast, 2 structures with binary effect appeared exclusively in the saline condition: RPA (Nucleus Raphe Pallidus), SCO (Subcommissural organ).

### Acute exposure to morphine induces rapid increases in c-Fos activity in regions implicated in addiction

We then determined the impacts of different durations of morphine exposure on brain region activation by comparing the respective morphine and saline groups at 1h and 4h post-administration (Fig. 5a). c-Fos positive cells increased compared to saline controls in both 1h and 4h groups. However, at 4h, the difference between morphine and saline brains was more pronounced with the average fold change for the whole brain increasing from 1.2 at 1h to 2.3 at 4h. Similarly, the number of brain structures displaying statistically significant differences from saline control increased from 12 structures at 1h (Fig. 4b, Table S4) to 34 structures (Fig. 4c, Table S5) at 4h. Non-overlapping binary effects were observed for both groups (Table S3, Fig. S3a).

**Figure 5.**
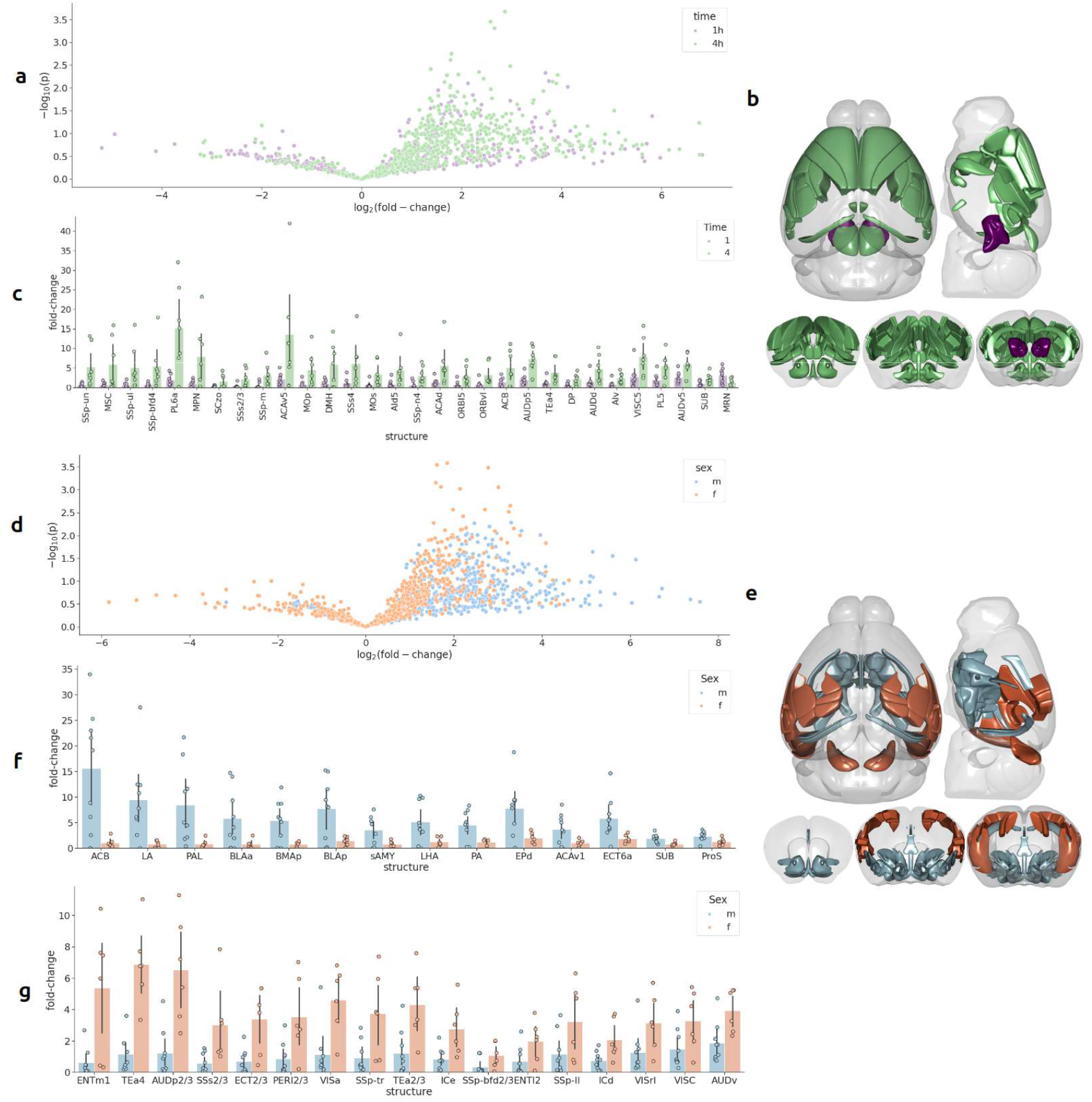
Time- and sex-dependent effects of morphine on region-specific brain activity. **(a)** The distribution of morphine over saline density fold-change by brain region at 1h (purple) and 4h (green) post-exposure as displayed in a volcano plot. **(b)** Brain regions where increased activity (p<0.05) was observed at 1h (purple) and 4h (green) are highlighted in a cartoon model of the brain. Density fold-change for each region is displayed in **(c)**. Bar plots are sorted (left->right) according to greatest fold-change. (**d)** The distribution of morphine over saline density fold-change by brain region for male (blue) and female (orange) animals is displayed in a volcano plot. (**e)** Brain regions where increased activity (p<0.05) was observed in male (blue) and female (orange) are highlighted in a cartoon model of the brain. Density fold-change for each region is displayed where activity is greater in (**f)** male and (**g)** female animals. Bar plots are sorted (left->right) according to greatest difference between male and female animals.

When we compared differences between 1h and 4h morphine groups, c-Fos density variations suggested dynamic changes in brain activity (Fig. S4a). Overall, 29 structures displayed statistically significant increases between 1h and 4h post administration (Fig. 5b). Whereas the Midbrain Reticular Nucleus (MRN) was the only structure to show a statistically significant decrease over this period at 2.49 fold (Fig. 5c). Many of the significant structures at 4h belonged to the cortex-basal ganglia loop^22,23^, reward/motivation^20,24^ and memory^25,26^ circuits including the Nucleus Accumbens (ACB), Prelimbic area (PL), Agranular Insular Cortex (AI)^25^, Orbitofrontal Cortex (ORB), Primary and Secondary motor areas (MOp, MOs), Anterior Cingulate Area (ACA) and Subiculum (SUB).

### Male and female mice display distinct patterns of c-Fos expression in response to morphine

We then examined whether the response to morphine differed based on sex. When we compared morphine and saline controls (Fig. 5d), activity was overall greater in male animals which displayed a 1.63 fold increase compared to female animals that displayed a 1.55 fold increase. The number of structures with statistically significant responses to morphine were similar between male and female groups with 32 structures responding in males (Fig. 4d, Table S7) and 31 structures responding in females (Fig. 4e, Table S6). Among those structures, 9 regions were seen in both sexes: Dorsal auditory area, layer 5 (AUDd5), Primary auditory area, layer 5 (AUDp5), AUDv5, Field CA3 in the Hippocampus, Entorhinal area, lateral part, layer 5 (ENTl5), Parabrachial nucleus (PB), Supplemental somatosensory area, layer 5 (SSs5), Visceral area, layer 5 (VISC5), Postrhinal area, layer 5 (VISpor5). The remaining structures that differed between sexes displayed distinct differences in the anatomical distribution (Fig. 4d-e, S4b) Notably, more structures displayed binary effects in male mice with 25 structures in contrast to female mice with only 1 binary structure (Table S3).

When we directly compared fold-change in morphine exposed mice between male and female groups, we identified 31 brain structures with significant differences between the sexes. Male mice displayed 14 structures with a greater fold-change when compared to females (Fig. 5f) whereas female mice demonstrated a greater fold change in the remaining 17 structures (Fig. 5g). Among the structures with more pronounced activity in male mice were members of the reward/motivation circuit including the Nucleus Accumbens (ACB), Pallidum (PAL) and Lateral Hypothalamus (LHA)^31^; Limbic system and the extended Amygdala (LA, BLAa, BLAp, BMAp, sAMY, PA, ECT)^32^; and the Memory circuits including Anterior Cingulate Area (ACA) and the Subiculum (SUB)^30,31^. In contrast, female mice showed more pronounced activity in structures associated with sensory inputs, association and memory, including visual areas (prefix: VIS), auditory areas (prefix: AUD), Inferior Colliculus (IC), Somatosensory areas (SS), Visceral areas (VISC), Entorhinal areas (ENT), Perirhinal areas (PERI), and Temporal Association areas (TEa).

## Discussion

In our study, we present a fast, high-throughput approach for mapping drug-evoked neuronal activity in the brains of mice acutely exposed to morphine. Our approach combined high speed microscopic imaging techniques, high-performance computing workflows and validated open-source software tools to identify and map cells within the whole mouse brain. We collected high-resolution whole brain data across sexes and times after morphine exposure. The datasets will be made publicly available following peer review and at the time of publication, along with all derived data, the code and computational environments used for analysis. Our workflow revealed time-, region-, and sex-dependent differences in c-Fos activity throughout discrete regions of the whole brain. Indeed, our results indicate dynamic subregion-specific changes in cell activity that became more pronounced 4h following drug exposure. This includes changes in the reward circuit^20^ and more broadly, cortex-basal ganglia loop that is involved in reinforcement learning and choice of behavioral programs^33^. Moreover, male and female animals displayed differences in their morphine responses with males displaying increased activity overall compared to females. This included higher activation in brain regions that belong to the mesolimbic dopamine system^34^, broader reward circuits^20^, and the extended amygdala^35^ implicated in addiction including Nucleus Accumbens, Basolateral Amygdala and Cingulate Cortex. While the mesolimbic dopamine system is activated more extensively in male mice, the association and sensory related cortical regions are activated more extensively in female mice. Male mice also displayed greater numbers of regions with binary effects where activity was only detected following exposure to morphine but not to saline. Collectively, our work opens the door to a robust, unbiased approach to identify novel drug-induced temporal and spatial changes in whole brain activity at cellular resolution.

We found region-specific neuronal activation in response to acute morphine treatment across 46 brain structures. Although our data confirmed activity in previously reported brain regions^36^, we also found unexpected morphine effects in regions not traditionally studied in the context of opioid actions including Raphe nuclei (dorsal, midbrain, pontine), Hippocampo-amygdalar transition area, and basolateral amygdala (BLA). Notably, our data are consistent with previous work showing that most of the structures also express mu opioid receptors (MOR) according to a brain atlas of MOR expression^37^. Interestingly, we identified strong morphine-induced activation of c-Fos expression in the BLA despite relatively weak MOR. We posit that this discrepancy may be due to an indirect circuit-level effect where upstream MOR activation may drive the activity of BLA cells downstream of the original morphine stimulation. Consistent with this, opioids modulate the pain circuitry both directly and indirectly via actions on serotonergic neurons that extend from the PAG through the rostral ventromedial medulla enroute to the spinal cord^38^. These potential MOR-serotonin links are also evident in the prominent morphine-induced activation of the DR and related Raphe nuclei, structures classically associated with the serotonin system, which is in line with previous research^38–42^.

We identified distinct sex differences in the response to morphine in male versus female mice, consistent with growing evidence of sex differences associated with opioid actions both clinically and in preclinical models^43,44^. In preclinical models, females demonstrate greater opioid self-administration compared to males^45–48^. Likewise, clinically, females are ∼4-fold more likely to inject heroin compared to males^49,50^. Interestingly, we found that males exhibited more extensive morphine-induced cell activation in the Striatum, Pallidum, Amygdala and Hippocampus while female mice showed significantly more c-Fos expression primarily in the Isocortex. This raises the possibility that these sex differences are due to sexually dimorphic patterns of MOR expression. Consistent with this, females possess a higher density of MOR-expressing neurons in several cortical regions including ACA and SS versus males^51^. It is also possible that these sex differences are at least partially mediated by estrogen, given estrogen’s emerging role as a modulator of the brain opioid system^49,52^. Future work will be required to elucidate the mechanisms underlying the interactions between opioid actions, sex, and brain regions.

We have also found temporal differences in cell activation at 1h versus 4h time points. At 1h following drug administration, there were fewer anatomic structures with significant differences between morphine and saline conditions. Behavioral studies have demonstrated onset of morphine-induced hyperlocomotion within 20-min of drug administration^53^, suggesting that 1h of exposure is sufficient for alterations in cell activity within select brain regions. However, by 4h post-administration, the data are less variable, with clearer cell activity within structures traditionally related to the actions of drugs of abuse including the Nucleus Accumbens, prelimbic area, infralimbic area, orbital area, motor area, agranular insular area and anterior cingulate area. Our findings are therefore in agreement with previous studies showing that c-Fos expression peaks in both morphine and saline brains at 30-60 minutes post-injection followed by a return to baseline in saline but not morphine conditions^54^.

We note that our use of c-Fos labeling to identify morphine-induced neuronal activity may capture the strongest stimuli but still miss other groups of cells with less robust patterns of activation^55^. This can sometimes be exacerbated by the inherent properties of the brain tissue and imperfections in clearing which lead to light absorption and scattering, ultimately reducing signal-to-noise deep within the tissue. To increase our chances of detecting low signal-to-noise events, our workflow leveraged two methods of cell detection combining conventional threshold-based detection and neural networks that are threshold independent. While c-Fos detection acknowledges activity-driven expression, this approach does not permit measurement of the magnitude of cell activation. Thus, we cannot use our approach to quantitatively ascertain whether already-activated cells exhibit change of expression in response to different durations of morphine exposure.

Our analysis did not make conclusions about structures in the CCFv3 below a 0.25mm^3^ volume threshold which included 270 structures within the grey matter. This was a deliberate choice designed to reduce any inaccuracies in our analysis that might have been introduced due to errors in the alignment to the CCFv3 template. Very small brain regions require exceedingly accurate alignment. Inaccuracies in the multimodal alignment of large volumetric datasets are to be expected. Modality-specific atlas reference images^56^ exist for other imaging modalities and might improve accuracy, but currently there is no confocal-specific template for the CCFv3. We expect that small inconsistencies in alignments would average out with greater numbers of animals per group. Thus, we expect that future studies will focus on imaging a greater number of animals which will enable us to obtain both increased accuracy and statistical power.

### Conclusions

Recent technological advances in tissue clearing and whole brain imaging have enabled us to digitally reconstruct entire mouse brains at cellular resolution. By merging high-speed RSCM^14^ and automated high-performance computing workflows designed to detect and map individual cells to locations within the brain, we can comprehensively map drug-activated neurons across dozens of animals in an unbiased manner at single-cell resolution. The resulting data demonstrate broad patterns of increased activity within the brains of morphine-exposed animals in a sex- and region-specific manner. Importantly, a single experiment allowed us to confirm the role of structures described in a diverse literature focused on specific brain regions and simultaneously identify novel structures which vary with time and sex. Overall, our fast whole brain imaging combined with massively parallel analysis opens avenues for exploring a large parameter space including actions of different drugs, different time courses of drug action, as well as sex- and region-specific effects. These studies therefore offer a more comprehensive understanding of the actions of opioids and offer targets for the development of novel therapeutic approaches.

## Methods

### Animals

Adult male and female mice (C57BL/6J, Jackson Laboratory, stock # 000664), (Total N=27; 15 males, 12 females, 8–10-weeks-old). All animals were housed in a 12/12 light cycle (7 am lights on, 7 pm lights off). Water and rodent chow were provided ad libitum throughout the experiment. Animals were habituated to the laboratory environment for 1 week prior to experimental testing. All procedures were approved and performed in compliance with the Institutional Animal Care and Use Committee at Boston University and the University of Pittsburgh.

### Drugs

All drugs were obtained from Sigma-Aldrich (St. Louis, MO) unless noted otherwise. Morphine sulfate was dissolved in a solution of filtered (0.22μm) 0.9% saline.

### Drug Treatment and Brain Collection

Mice were randomly assigned to 1h or 4h treatment groups. Animals were injected i.p. with either morphine (10mg/kg) or saline (10mg/kg, w/v). Morphine solution or saline were injected intraperitoneally (i.p.) in awake animals that were lightly restrained. Mice were immediately placed individually in a novel environment consisting of a plexiglass chamber with animal bedding. Animals were allowed to explore this environment for 1h before undergoing brain collection (1h group) or returned to their home cage for 3-hours before brain collection at the 4h time point (4h group). Animals underwent transcardiac perfusion with ice-cold PBS until blood had been cleared from the mouse. This was followed by 4% paraformaldehyde (PFA) perfusion for 10 minutes. Fixed brains were dissected and then transferred to 4% PFA at 4ºC for 24 hours before transferring to ice-cold PBS.

### Tissue Clearing and Staining

Brains were prepared using a mixture of CUBIC^57^, 3DISCO^58^, and SWITCH^59^ clearing and staining protocols. All steps took place at room temperature. The samples were pre-cleared using 50% CUBIC R1 for 1-2 weeks, changing the solution every 24-48 hours. After pre-clearing, CUBIC R1 was rinsed out and samples were stained with a rabbit monoclonal anti-phospho-c-Fos primary antibody (catalog# 8677S, Cell Signaling Technology, Danvers, MA) according to the SWITCH protocol. Samples were equilibrated in SWITCH-Off solution (5 mM SDS + 0.04% NaAz) for 24h, followed by addition of the anti-c-Fos primary antibody (Rb) to the samples (1:250 in SWITCH-Off solution). Samples incubated in primary antibody for 1 week, then washed for 24-48 hours in SWITCH-On solution (1X PBS + 0.04% NaAz + TritonX). This process was repeated with Cy5-conjugated goat and donkey anti-rabbit secondary antibodies (1:2000, catalog#s 111-605-003 and 711-175-152, Jackson Immunoresearch, West Grove, PA) in SWITCH-Off. Samples were post-fixed in 4% paraformaldehyde for 1h. After staining, samples were dehydrated in 2-hour cycles of increasing concentrations of tetrahydrofuran (%THF: 30, 50, 70, 90 (x2), 100 (x2)). Following dehydration, samples were cleared and mounted in dibenzyl ether to be imaged.

### Imaging

Brains were imaged via ribbon scanning confocal microscopy (RSCM, CALIBER I.D. RS-G4, Andover, MA, USA) as described previously^14^. Briefly, up to 8 brains were immersed in dibenzyl ether and arrayed in a single imaging chamber. The chamber was sealed with a glass coverslip and then filled with glycerol. We used a Nikon 20x, 1.0NA, 8.2mm WD, glycerol immersion (CF190) objective specifically designed for cleared tissue imaging. Data were acquired using a 488 (autofluorescence filter Semrock FF01-520/44 filter) and 647 (Cy5 c-Fos signal - filter Semrock FF01-670/30) filter with full sequencing enabled. Four samples were imaged at 0.337μm x 0.337μm x 4.57μm (X,Y,Z) voxel resolution, and twenty-three samples were imaged with 0.361μm x 0.361μm x 5.33μm (X,Y,Z) voxel resolution. The data were stitched and assembled into 2D images each representing a single channel and z-plane. Shot noise was then removed from the images using a convolutional neural network based on noise-2-noise^60^ and trained on RSCM data. Image planes were then assembled into a 3D volume using the freely available Imaris File Converter (Bitplane), resulting in a single Imaris file composed of a multiscale chunked data structure that represented the entire whole brain dataset. All subsequent downstream analysis was completed from these files.

### Compute

We produced an automated c-Fos mapping pipeline which included the detection of c-Fos-positive cells and the alignment of each brain to the Allen Mouse Brain Common Coordinate Framework v3^17^ (CCFv3). Image processing was performed using the Center for Biologic Imaging’s high performance computing environment. The environment is schedulable via SLURM^18^ and consists of a 18 node compute partition having 24 cores and 96GB RAM and a 5 node GPU partition each with 72 cores, 1.5TB RAM and 8x NVIDIA P40. All nodes mount two BeeGFS high performance file systems consisting of 7PB of HDD and 160TB of SSD. RSCM data was acquired directly as unstitched ribbons to the BeeGFS SSD storage system prior to being stitched and assembled on the compute nodes as 2D images. Data was then copied to the HDD storage system where they were queued for denoising on the GPU nodes. Following denoising, all images were assembled into Imaris (.ims) files which were the basis for all remaining image analysis. The final size of each full brain dataset was on average ∼1.5 terabytes, comprising 40TB across 27 brains. During image generation and processing, these data were transformed 3 times transiently generating approximately 200TB. The final denoised, ims files were retained and are publicly available at the Brain Image Library as discussed below. Data mining and analysis was developed and tested on custom built workstations consisting of 32-core, 64-thread AMD Ryzen Threadripper PRO, 256GB RAM, and NVIDIA A6000 GPU.

### c-Fos puncta detection

Data contained fluorescent c-Fos-positive puncta of different sizes and intensities. We found that the larger and brighter spots were well detected with cellfinder^61^ python tool, whereas small and dim spots were detectable with the deepblink particle model^62^ based on UNet architecture^63^. Both algorithms were applied to each imaged brain. Together, these algorithms were purposely biased towards over detection to ensure that all cells were identified. Thus, to avoid duplicate detections, the resulting spots were run through DBSCAN clustering algorithm^64^ which clustered the duplicated detections together and replaced these clusters by a single point. In addition, detected points contained artifacts such as edges, noise peaks and autofluorescent vasculature. To eliminate these artifacts, each detected spot was fed to a deep learning binary classifier based on ResNet-50^65^. The model was trained on >40,000 cell and non-cell examples across multiple brains from this study. To confirm precision, recall, and f1^66^ score of the detection and classification process, ground truth data was annotated manually in a small patch of size of 25 × 250 × 250 pixels from each brain. For brains where the f1 score was <80%, additional improvement of classification was needed. For this, the model was fine-tuned on a relatively small number (50-200 examples of cells and non-cells) of manually annotated data from each of these brains, using fastai^67^. Finally, the data was inspected visually, and for a few brains where classification still did not reach a f1 score of ≥80%, minor manual removal of residual artifacts was done. Optionally, removal of background detections can be done to reduce the time spent on clustering duplicated detections and cell/artifact classification. We created background-foreground masks for each brain using accelerated pixel and object classification plugin for napari^68^ that employs random forest machine learning algorithm^69^. These masks will be provided alongside the raw data.

### Registration to the CCFv3

Each brain was registered to the 10μm CCFv3 following downsampling of the full resolution to 10 μm isotropic resolution. Brainreg^70^ a fully automated 3D registration python package was used for all registrations. The registration results were inspected visually. If the registration quality was not satisfactory, the raw data was pre-processed by filtering in the frequency domain, contrast stretching, and background subtraction until a visually good alignment was obtained. The resulting deformation field was used to transform each of the c-Fos points into CCFv3 space. Every c-Fos-positive cell was associated with an Allen Mouse Brain Atlas parcellation where it was located.

Critically, tissue clearing results in some physical deformation of the mouse brain. Although relatively rare, the stresses placed on delicate brain tissue can sometimes cause tearing or result in tissue loss. Though nonlinear alignment algorithms are used in brainreg as an attempt to correct for changes in tissue shape, they are never entirely accurate. Thus, to reduce errors resulting from minor inaccuracies in alignment, we eliminated the smallest brain structures by focusing our downstream analysis on structures that were at least 0.25 mm^3^. At this scale, we expect that inaccuracies in alignment will be compensated for by averaging data from multiple brains.

### Data analysis

Our workflow produced a CSV table, which included the locations of c-Fos-positive cells mapped to the CCFv3 and labeled with the associated parcellation. These secondary data tables will be made public alongside the raw data as mentioned below. For analysis, tables for all of the brains were combined and used for data mining. Each row in a corresponding table contained the coordinates for a spot in the raw brain image (world space), coordinates for a spot the CCFv3, and the spot’s corresponding CCFv3 parcellation (*i*.*e*., brain structure). CSV files for all brains were loaded into an interactive Jupyter^71^ notebook for analysis. For each brain, the points were aggregated by brain structure, and the density of c-Fos-positive cells for each structure was calculated as the number of spots in the structure over the volume of the structure. Densities were the main metric used to make comparisons of structures across brains from different animals and different experimental conditions.

### Visualizations

Raw data was stored in the Imaris file format, and Imaris viewer was used for initial visual inspection. Volumetric brain cartoons were made using brainrender^72^. Bar and pie charts were done using Plotly^73^ or Seaborn^74^. Heatmaps in the CCFv3 were made in napari^75^.

### Statistical analysis

To calculate p-values between different experimental conditions in bulk, in an automatic way, for all brain structures, we used student’s 2-sample t-test, available through scipy^76^. Numbers of animals for each group ranged from 2-5. For the analysis, we pooled statistically independent groups together, therefore the n was always ≥5. Unless otherwise indicated, data is represented as mean ± standard error of the mean.

## Data Availability

At the time of publication, all full resolution imaging data described herein will be deposited in the Brain Image Library (BIL) and be made available along with the derived analysis data under an associated DOI.

## Code Availability

All code used to generate the results described in this manuscript will be deposited in github and/or made available alongside the imaging data deposited at BIL. Jupyter notebooks will be provided which reproduce the figures and tables.

## Acknowledgements

We are grateful for the discussions and technical assistance provided by Dr. Sam Golden. This study was supported by the National Institutes of Health (R01DK124219 to Z.F.; R01ES034037 to Z.F., R36DA057972 to J.K.; T32GM133353 to J.K., R01DA061243 to R.W.L. and Z.F., R21DA052419 to R.W.L., Z.F., and A.M.W., R24MH114793 to A.M.W.), the Department of Defense (PR210207 to Z.F.), the Chan Zuckerberg Initiative DAF (2023-329680 to K.M.K) and the Lundbeck Foundation (R276-2018-792 and R359-2020-2301 to U.G.).

## Author contributions statement

I.V., M.C.S., S.P., J.K., P.N.J., A.M.W. conducted the experiments. I.V., M.C.S, E.A., K.H., T.L., K.M.K., J.J.S., Z.F., M.D.L., A.M.W. analyzed the results. I.V., M.C.S., Z.F., A.M.W wrote the manuscript. R.W.L., Z.F., A.M.W. conceived the experiment(s). U.G., R.W.L., Z.F., A.M.W. supervised the work, and contributed funding. All authors reviewed the manuscript.

## Abbreviations

BIL: Brain Image Library
CCFv3: Allen Brain Atlas Common Coordinate Framework Version 3
CSV: Character Separated Values
FAIR: Findable, Accessible, Interoperable, Reusable
GPU: Graphical Processing Unit
HDD: Hard Disk Drive
IEG: Immediate Early Gene
i.p: intraperitoneal
MOR: Mu Opioid Receptor
OUD: opioid use disorder
PB: petabyte
RAM: Random Access Memory
RSCM: Ribbon Scanning Confocal Microscopy
SLURM: Simple Linux Utility for Resource Management
SSD: Solid State Drive
TB: Terabyte

## Brain Regions found within the CCFv3

AAA: Anterior Amygdalar Area
ACA: Anterior Cingulate Area
ACAd1: Anterior Cingulate Area, dorsal part, layer 1
ACAd2/3: Anterior Cingulate Area, dorsal part, layers 2/3
ACAd5: Anterior Cingulate Area, dorsal part, layer 5
ACAd6a: Anterior Cingulate Area, dorsal part, layer 6a
ACAv1: Anterior Cingulate Area, ventral part, layer 1
ACAv2/3: Anterior Cingulate Area, ventral part, layers 2/3
ACAv5: Anterior Cingulate Area, ventral part, layer 5
ACAv6a: Anterior Cingulate Area, ventral part, layer 6a
ACB: Nucleus Accumbens
AId: Agranular insular area, dorsal part
AId1: Agranular Insular Area, dorsal part, layer 1
AId2/3: Agranular Insular Area, dorsal part, layers 2/3
AId5: Agranular Insular Area, dorsal part, layer 5
AId6a: Agranular Insular Area, dorsal part, layer 6a
AI: Agranular Insular Area
AIp1: Agranular Insular Area, posterior part, layer 1
AIp2/3: Agranular Insular Area, posterior part, layers 2/3
AIp5: Agranular Insular Area, posterior part, layer 5
AIp6a: Agranular Insular Area, posterior part, layer 6a
AIv2/3: Agranular Insular Area, ventral part, layers 2/3
AIv5: Agranular Insular Area, ventral part, layer 5
AHN: Anterior Hypothalamic Nucleus
AOBmi: Accessory Olfactory Bulb, mitral layer
AON: Anterior Olfactory Nucleus
APN: Anterior Pretectal Nucleus
APr: Area Prostriata
ARH: Arcuate Hypothalamic Nucleus
AM: Anteromedial Nucleus
AV: Anteroventral Nucleus
BLA: Basolateral amygdalar nucleus
BLAa: Basolateral Amygdalar Nucleus, anterior part
BLAp: Basolateral Amygdalar Nucleus, posterior part
BLAv: Basolateral Amygdalar Nucleus, ventral part
BMA: Basomedial amygdalar nucleus
BMAa: Basomedial Amygdalar Nucleus, anterior part
BMAp: Basomedial Amygdalar Nucleus, posterior part
BST, BNST: Bed nuclei of the stria terminalis
CA1: Cornu Ammonis area 1
CA2: Cornu Ammonis area 2
CA3: Cornu Ammonis area 3
COA: Cortical amygdalar area
COAa: Cortical Amygdalar Area, anterior part
COApl: Cortical Amygdalar Area, posterolateral part
COApm: Cortical Amygdalar Area, posteromedial part
CP: Caudoputamen
CTXsp: Cortical Subplate
DG-mo: Dentate Gyrus, molecular layer
DG-po: Dentate Gyrus, polymorphic layer
DG-sg: Dentate Gyrus, granule cell layer
DMH: Dorsomedial Hypothalamic Nucleus
DMX: Dorsal motor nucleus of the vagus nerve
DP: Dorsal Peduncular Area
DCO: Dorsal Cochlear Nucleus
DR: Dorsal Raphe Nucleus
ECT: Ectorhinal Area
ECT1: Ectorhinal Area, layer 1
ECT2/3: Ectorhinal Area, layers 2/3
ECT5: Ectorhinal Area, layer 5
ECT6a: Ectorhinal Area, layer 6a
EP: Endopiriform nucleus
EPd: Endopiriform Nucleus, dorsal part
EPv: Endopiriform Nucleus, ventral part
ENT: Entorhinal Area
ENTl1: Lateral Entorhinal Area, layer 1
ENTl2: Lateral Entorhinal Area, layer 2
ENTl3: Lateral Entorhinal Area, layer 3
ENTl5: Lateral Entorhinal Area, layer 5
ENTl5: Lateral Entorhinal Area, layer 5
ENTl6a: Lateral Entorhinal Area, layer 6a
ENTm1: Medial Entorhinal Area, layer 1
ENTm2: Medial Entorhinal Area, layer 2
ENTm3: Medial Entorhinal Area, layer 3
ENTm5: Medial Entorhinal Area, layer 5
ENTm6: Medial Entorhinal Area, layer 6
FRP: Frontal pole, cerebral cortex
FRP1: Frontal Pole, layer 1
FRP5: Frontal Pole, layer 5
FS: Fundus of Striatum
GU: Gustatory areas
GU2/3: Gustatory Area, layers 2/3
GU5: Gustatory Area, layer 5
GU6a: Gustatory Area, layer 6a
GPe: Globus Pallidus, external segment
GPi: Globus Pallidus, internal segment
GRN: Gracile Nucleus
HPF: Hippocampal Formation
HATA: Hippocampus-Amygdala Transition Area
HY: Hypothalamus
ILA: Infralimbic Area
IC: Inferior Colliculus
ICc: Inferior Colliculus, central nucleus
ICd: Inferior Colliculus, dorsal cortex
ICe: Inferior Colliculus, external cortex
IO: Inferior Olive
IPN: Interpeduncular Nucleus
IRN: Intermediate Reticular Nucleus
LA: Lateral Amygdalar Nucleus
LAV: Lateral Vestibular Nucleus
LD: Lateral Dorsal Nucleus
LHA: Lateral Hypothalamic Area
LGd-co: Lateral Geniculate Nucleus, dorsal, core region
LGv: Lateral Geniculate Nucleus, ventral part
LP: Lateral Posterior Nucleus
LS: Lateral septal nucleus
LSc: Lateral Septum, caudal part
LSr: Lateral Septum, rostral part
LSv: Lateral Septum, ventral part
LPO: Lateral preoptic area
LRNm: Lateral Reticular Nucleus, medial part
MA: Medial Amygdala
MEA: Medial Amygdalar Nucleus
MB: Midbrain
MGm: Medial Geniculate Nucleus, magnocellular division
MGv: Medial Geniculate Nucleus, ventral division
MM: Medial Mammillary Nucleus
MO: Somatomotor areas
MOB: Main Olfactory Bulb
MOp: Primary Motor Area
MOp: Primary motor area
MOp1: Primary Motor Area, layer 1
MOp2/3: Primary Motor Area, layers 2/3
MOp5: Primary Motor Area, layer 5
MOp6a: Primary Motor Area, layer 6a
MOs: Secondary Motor Area
MOs1: Secondary Motor Area, layer 1
MOs2/3: Secondary Motor Area, layers 2/3
MOs5: Secondary Motor Area, layer 5
MOs6a: Secondary Motor Area, layer 6a
MARN: Magnocellular Reticular Nucleus
MDRN: Medullary Reticular Nucleus
MS: Medial Septum
MRN: Midbrain Reticular Nucleus
MY: Medulla
NAc: Nucleus Accumbens
NDB: Nucleus of the Diagonal Band
NPC: Nucleus of the Posterior Commissure
NLOT: Nucleus of the lateral olfactory tract
NLL: Nucleus of the lateral lemniscus
NTS: Nucleus of the Solitary Tract
ORB: Orbital Area
ORBl: Orbital area, lateral part
ORBl1: Orbital Area, lateral part, layer 1
ORBl2/3: Orbital Area, lateral part, layers 2/3
ORBl5: Orbital Area, lateral part, layer 5
ORBl6a: Orbital Area, lateral part, layer 6a
ORBm: Orbital area, medial part
ORBm1: Orbital Area, medial part, layer 1
ORBm2/3: Orbital Area, medial part, layers 2/3
ORBm5: Orbital Area, medial part, layer 5
ORBvl: Orbital area, ventrolateral part
ORBvl1: Orbital Area, ventrolateral part, layer 1
ORBvl2/3: Orbital Area, ventrolateral part, layers 2/3
ORBvl5: Orbital Area, ventrolateral part, layer 5
OT: Olfactory Tubercle
PA: Posterior Amygdalar Nucleus
PAA: Piriform-Amygdalar Area
PAL: Pallidum
PAG: Periaqueductal Gray
PB: Parabrachial Nucleus
p5: Pontine Reticular Formation, p5 Subregion
PCG: Pontine Central Gray
PG: Pontine Gray
PIR: Piriform area
POR: Pontine Reticular Formation, Oral Part
PRNc: Pontine Reticular Nucleus, Caudal Part
PRNr: Pontine Reticular Nucleus, Rostral Part
PS: Parastrial nucleus
PSV: Pontine Superior Vestibular Nucleus
PVT: Paraventricular nucleus of the thalamus
RAmb: Nucleus ambiguus, rostral part
RPA: Raphe Pallidus
RE: Nucleus of reuniens
RPO: Raphe Pontis Nucleus
RO: Raphe Obscurus Nucleus
RSP: Retrosplenial area
RSPagl: Retrosplenial area, lateral agranular part
RSPagl1: Retrosplenial Area, agranular, lateral part, layer 1
RSPagl2/3: Retrosplenial Area, agranular, lateral part, layers 2/3
RSPagl5: Retrosplenial Area, agranular, lateral part, layer 5
RSPagl6a: Retrosplenial Area, agranular, lateral part, layer 6a
RSPd: Retrosplenial area, dorsal part
RSPd1: Retrosplenial Area, dorsal part, layer 1
RSPd2/3: Retrosplenial Area, dorsal part, layers 2/3
RSPd5: Retrosplenial Area, dorsal part, layer 5
RSPd6a: Retrosplenial Area, dorsal part, layer 6a
RSPv: Retrosplenial area, ventral part
RSPv1: Retrosplenial Area, ventral part, layer 1
RSPv2/3: Retrosplenial Area, ventral part, layers 2/3
RSPv5: Retrosplenial Area, ventral part, layer 5
RSPv6a: Retrosplenial Area, ventral part, layer 6a
RN: Red Nucleus
SCO: Subcommissural Organ
SC: Superior colliculus
SCdg: Superior Colliculus, deep gray layer
SCdw: Superior Colliculus, deep white layer
SCig: Superior Colliculus, intermediate gray layer
SCiw: Superior Colliculus, intermediate white layer
SCop: Superior Colliculus, optic layer
SCsg: Superior Colliculus, superficial gray layer
SCzo: Superior Colliculus, zonal layer
SF: Septofimbrial Nucleus
SOCl: Superior Olivary Complex, Lateral Part
SPIV: Spinal Trigeminal Nucleus, ventral part
SPVC: Spinal Trigeminal Nucleus, caudal part
SPVI: Spinal Trigeminal Nucleus, intermediate part
SPVO: Spinal Trigeminal Nucleus, oral part
SS: Somatosensory Area
SSP-bfd1: Primary Somatosensory Area, Barrel Field, layer 1
SSP-bfd2/3: Primary Somatosensory Area, Barrel Field, layers 2/3
SSP-bfd4: Primary Somatosensory Area, Barrel Field, layer 4
SSP-bfd5: Primary Somatosensory Area, Barrel Field, layer 5
SSP-bfd6a: Primary Somatosensory Area, Barrel Field, layer 6a
SSp-bfd2/3: Primary Somatosensory Area, Barrel Field, layers 2/3
SSp-ll: Primary Somatosensory Area, Lower Limb
SSp-ll1: Primary Somatosensory Area, Lower Limb, layer 1
SSp-ll2/3: Primary Somatosensory Area, Lower Limb, layers 2/3
SSp-ll4: Primary Somatosensory Area, Lower Limb, layer 4
SSp-ll5: Primary Somatosensory Area, Lower Limb, layer 5
SSp-ll6a: Primary Somatosensory Area, Lower Limb, layer 6a
SSp-m: Primary Somatosensory Area, Mouth
SSp-m1: Primary Somatosensory Area, Mouth, layer 1
SSp-m2/3: Primary Somatosensory Area, Mouth, layers 2/3
SSp-m4: Primary Somatosensory Area, Mouth, layer 4
SSp-m5: Primary Somatosensory Area, Mouth, layer 5
SSp-m6a: Primary Somatosensory Area, Mouth, layer 6a
SSp-n: Primary Somatosensory Area, Nose
SSp-n1: Primary Somatosensory Area, Nose, layer 1
SSp-n2/3: Primary Somatosensory Area, Nose, layers 2/3
SSp-n4: Primary Somatosensory Area, Nose, layer 4
SSp-n5: Primary Somatosensory Area, Nose, layer 5
SSp-n6a: Primary Somatosensory Area, Nose, layer 6a
SSp-tr: Primary Somatosensory Area, Trunk
SSp-tr2/3: Primary Somatosensory Area, Trunk, layers 2/3
SSp-tr5: Primary Somatosensory Area, Trunk, layer 5
SSp-ul: Primary Somatosensory Area, Upper Limb
SSp-ul1: Primary Somatosensory Area, Upper Limb, layer 1
SSp-ul2/3: Primary Somatosensory Area, Upper Limb, layers 2/3
SSp-ul4: Primary Somatosensory Area, Upper Limb, layer 4
SSp-ul5: Primary Somatosensory Area, Upper Limb, layer 5
SSp-ul6a: Primary Somatosensory Area, Upper Limb, layer 6a
SSp-un: Primary Somatosensory Area, Unassigned Region
SSp-un2/3: Primary Somatosensory Area, Unassigned Region, layers 2/3
SSp-un5: Primary Somatosensory Area, Unassigned Region, layer 5
SSp-un6a: Primary Somatosensory Area, Unassigned Region, layer 6a
SSs1: Secondary Somatosensory Area, layer 1
SSs2/3: Secondary Somatosensory Area, layers 2/3
SSs4: Secondary Somatosensory Area, layer 4
SSs5: Secondary Somatosensory Area, layer 5
SSs6a: Secondary Somatosensory Area, layer 6a
SUT: Supratrigeminal Nucleus
SUM: Supramammillary Nucleus
TEa: Temporal Association Area
TEa1: Temporal Association Area, layer 1
TEa2/3: Temporal Association Area, layers 2/3
TEa4: Temporal Association Area, layer 4
TEa5: Temporal Association Area, layer 5
TEa6a: Temporal Association Area, layer 6a
TR: Taenia Tecta, rostral part
TRN: Tegmental Reticular Nucleus
TRS: Triangular Septal Nucleus
TU: Tuberal Nucleus
UVU: Uvula
V: Vestibular Nucleus
VISC: Visceral Area
VISC1: Visceral Area, layer 1
VISC2/3: Visceral Area, layers 2/3
VISC4: Visceral Area, layer 4
VISC5: Visceral Area, layer 5
VISC6a: Visceral Area, layer 6a
VIS: Visual Area
VISa: Anterior area
VISa2/3: Visual Association Area, layers 2/3
VISa5: Visual Association Area, layer 5
VISal: Anterolateral Visual Area
VISam: Anteromedial Visual Area
VISl: Lateral visual area
VISl2/3: Lateral Visual Area, layers 2/3
VISl5: Lateral Visual Area, layer 5
VISli: Laterointermediate Visual Area
VISp: Primary visual area
VISp1: Primary Visual Area, layer 1
VISp2/3: Primary Visual Area, layers 2/3
VISp4: Primary Visual Area, layer 4
VISp5: Primary Visual Area, layer 5
VISp6a: Primary Visual Area, layer 6a
VISpl: Posterolateral Visual Area
VISpm: Posteromedial Visual Area
VISpm2/3: Posteromedial Visual Area, layers 2/3
VISpm5: Posteromedial Visual Area, layer 5
VISpor: Postrhinal visual area
VISpor2/3: Postrhinal (Visual) Area, layers 2/3
VISpor5: Postrhinal (Visual) Area, layer 5
VTA: Ventral Tegmental Area
VM: Ventromedial Nucleus
VPL: Ventral Posterolateral Nucleus
VPM: Ventral Posteromedial Nucleus
VAL: Ventral Anterior-Lateral Nucleus
XII: Hypoglossal Nucleus
ZI: Zona Incerta

## Supplemental Materials

**Figure S1.**
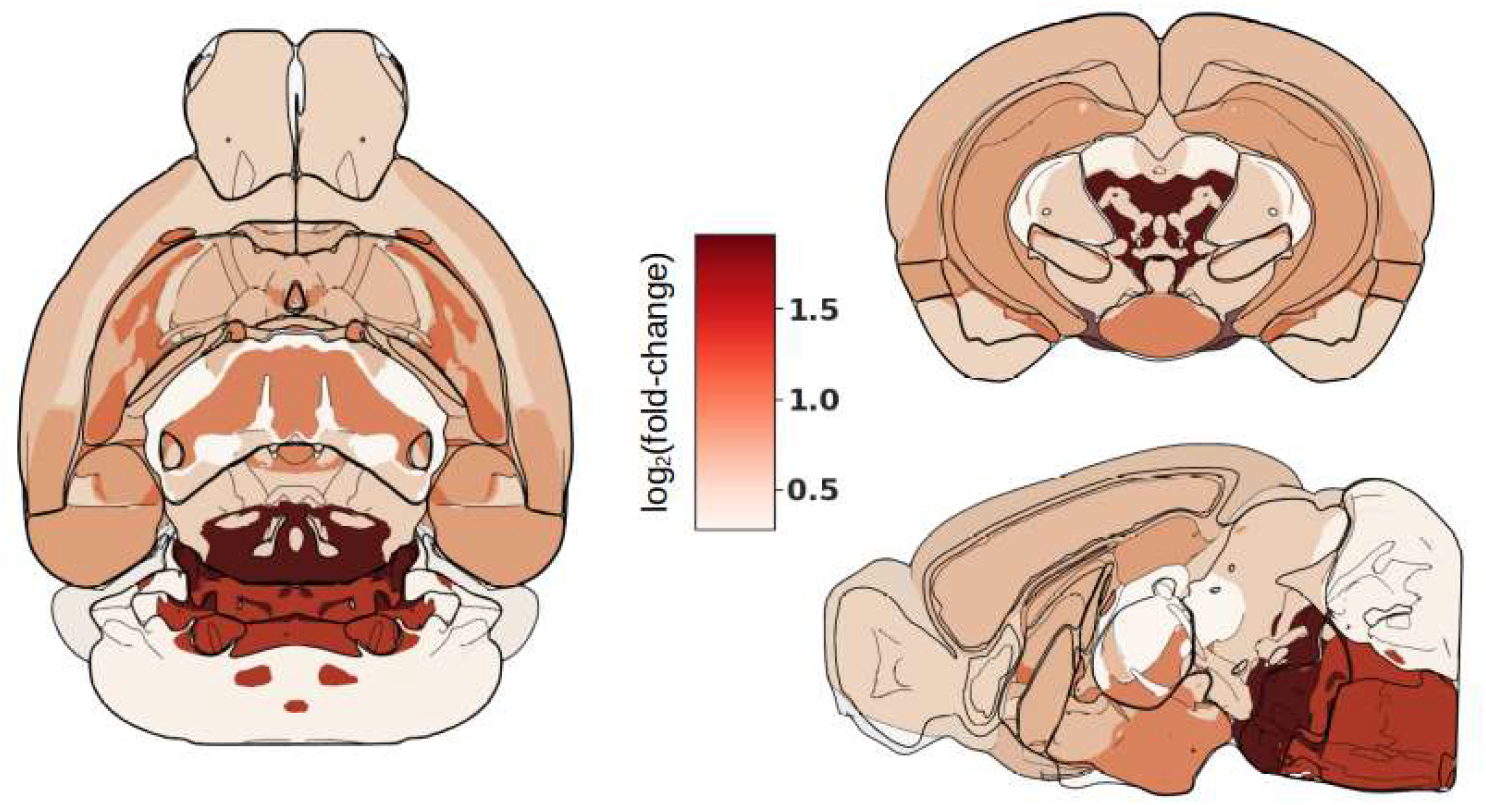
Morphine effect across large brain structures. A cartoon representation of the mouse brain with a heat map which displays the average density fold-change across structures summarized in Table 1. Average morphine cell densities are calculated across all 15 morphine treated brains in the study. Average saline densities are calculated across all 12 saline treated brains.

**Table S1.**
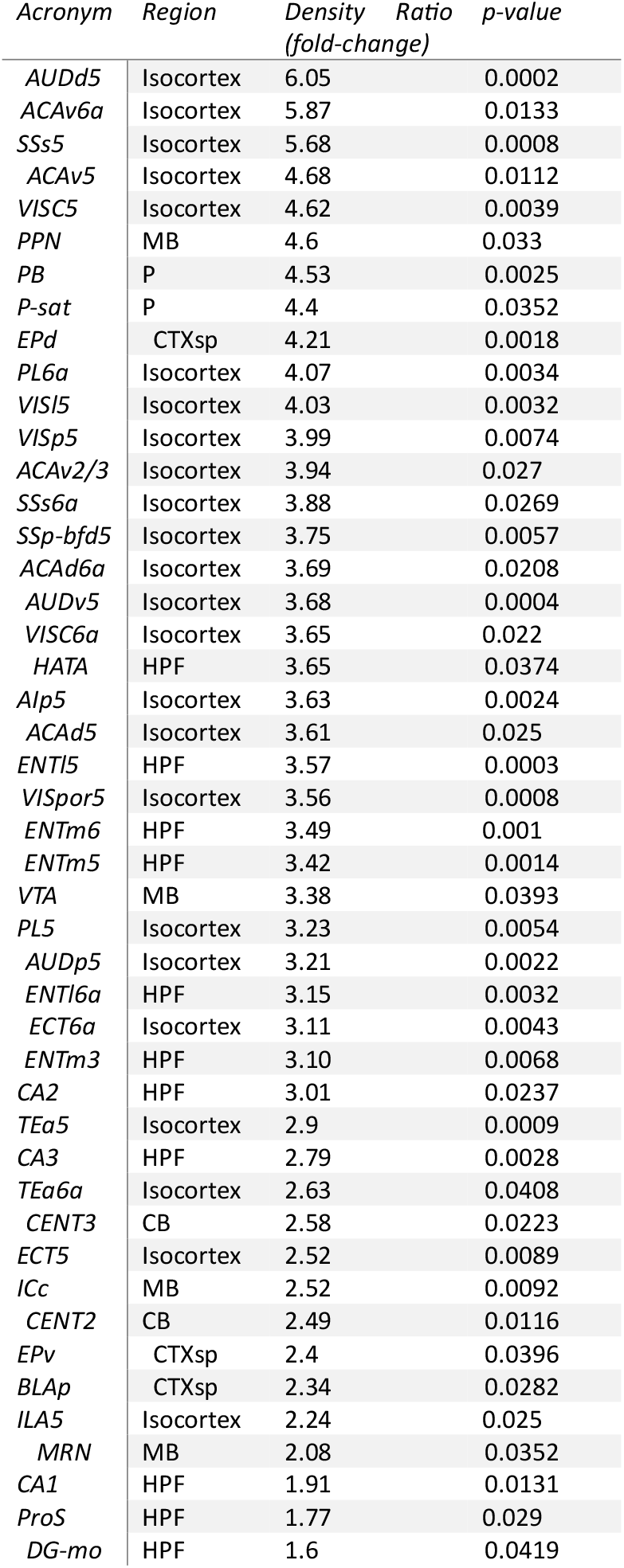
Allen Mouse Brain Atlas structures that exhibited a significant morphine effect. Table summarizing gray matter structures with significant morphine-induced changes in c-Fos expression (p<0.05), which were sufficiently large (>0.25 mm^3^ volume, to reduce effect of registration imperfections) and had at least 25 spots detected on average in both morphine and saline brains (to ensure reliable c-Fos puncta identification). 46 structures satisfied these conditions.

| *Acronym* | *Region* | *Density Ratio*<br>*(fold-change)* | *p-value* |
| --- | --- | --- | --- |
| *AUDd5* | Isocortex | 6.05 | 0.0002 |
| *ACAv6a* | Isocortex | 5.87 | 0.0133 |
| *SSs5* | Isocortex | 5.68 | 0.0008 |
| *ACAv5* | Isocortex | 4.68 | 0.0112 |
| *VISC5* | Isocortex | 4.62 | 0.0039 |
| *PPN* | MB | 4.6 | 0.033 |
| *PB* | P | 4.53 | 0.0025 |
| *P-sat* | P | 4.4 | 0.0352 |
| *EPd* | CTXsp | 4.21 | 0.0018 |
| *PL6a* | Isocortex | 4.07 | 0.0034 |
| *VISI5* | Isocortex | 4.03 | 0.0032 |
| *VISp5* | Isocortex | 3.99 | 0.0074 |
| *ACAv2/3* | Isocortex | 3.94 | 0.027 |
| *SSs6a* | Isocortex | 3.88 | 0.0269 |
| *SSp-bfd5* | Isocortex | 3.75 | 0.0057 |
| *ACAd6a* | Isocortex | 3.69 | 0.0208 |
| *AUDv5* | Isocortex | 3.68 | 0.0004 |
| *VISC6a* | Isocortex | 3.65 | 0.022 |
| *HATA* | HPF | 3.65 | 0.0374 |
| *Alp5* | Isocortex | 3.63 | 0.0024 |
| *ACAd5* | Isocortex | 3.61 | 0.025 |
| *ENTI5* | HPF | 3.57 | 0.0003 |
| *VISpor5* | Isocortex | 3.56 | 0.0008 |
| *ENTm6* | HPF | 3.49 | 0.001 |
| *ENTm5* | HPF | 3.42 | 0.0014 |
| *VTA* | MB | 3.38 | 0.0393 |
| *PL5* | Isocortex | 3.23 | 0.0054 |
| *AUDp5* | Isocortex | 3.21 | 0.0022 |
| *ENTl6a* | HPF | 3.15 | 0.0032 |
| *ECT6a* | Isocortex | 3.11 | 0.0043 |
| *ENTm3* | HPF | 3.10 | 0.0068 |
| *CA2* | HPF | 3.01 | 0.0237 |
| *TEa5* | Isocortex | 2.9 | 0.0009 |
| *CA3* | HPF | 2.79 | 0.0028 |
| *TEa6a* | Isocortex | 2.63 | 0.0408 |
| *CENT3* | CB | 2.58 | 0.0223 |
| *ECT5* | Isocortex | 2.52 | 0.0089 |
| *ICc* | MB | 2.52 | 0.0092 |
| *CENT2* | CB | 2.49 | 0.0116 |
| *EPv* | CTXsp | 2.4 | 0.0396 |
| *BLAp* | CTXsp | 2.34 | 0.0282 |
| *ILA5* | Isocortex | 2.24 | 0.025 |
| *MRN* | MB | 2.08 | 0.0352 |
| *CA1* | HPF | 1.91 | 0.0131 |
| *ProS* | HPF | 1.77 | 0.029 |
| *DG-mo* | HPF | 1.6 | 0.0419 |

**Table S2.** Allen Mouse Brain Atlas structures that trended towards significant morphine-induced effects. All CCFv3 structures that had morphine effects approaching significance (0.05 <= p < 0.1), contained gray matter, were sufficiently large (>0.25 mm^3^ volume, to reduce effect of registration imperfections and had at least 25 spots detected on average in both morphine and saline brains.

| *Acronym* | *Region* | *Density*<br>*Ratio (fold-*<br>*change)* | *p-value* |
| --- | --- | --- | --- |
| *CLA* | CTXsp | 9.32 | 0.071 |
| *Alp6a* | Isocortex | 5.82 | 0.066 |
| *Ald6a* | Isocortex | 4.90 | 0.051 |
| *GU5* | Isocortex | 4.24 | 0.057 |
| *RSPagl6a* | Isocortex | 4.14 | 0.066 |
| *MV* | MY | 3.99 | 0.097 |
| *MOp5* | Isocortex | 3.93 | 0.087 |
| *MOs6a* | Isocortex | 3.72 | 0.089 |
| *LPO* | HY | 3.70 | 0.053 |
| *MY-sen* | MY | 3.51 | 0.068 |
| *P-mot* | P | 3.44 | 0.071 |
| *RAmb* | MB | 3.37 | 0.06 |
| *SSp-m5* | Isocortex | 3.36 | 0.085 |
| *VISp6a* | Isocortex | 3.32 | 0.057 |
| *ORBl6a* | Isocortex | 3.15 | 0.091 |
| *SSp-bfd6a* | Isocortex | 2.87 | 0.06 |
| *MPN* | HY | 2.85 | 0.093 |
| *MPO* | HY | 2.77 | 0.065 |
| *AHN* | HY | 2.69 | 0.06 |
| *AAA* | STR | 2.57 | 0.073 |
| *PL2/3* | Isocortex | 2.5 | 0.082 |
| *Alv5* | Isocortex | 2.45 | 0.055 |
| *BLAv* | CTXsp | 2.42 | 0.065 |
| *SI* | PAL | 2.41 | 0.077 |
| *SCiw* | MB | 2.39 | 0.058 |
| *ACB* | STR | 2.34 | 0.072 |
| *VISli* | Isocortex | 2.31 | 0.081 |
| *ACAv1* | Isocortex | 2.29 | 0.061 |
| *AUDp* | Isocortex | 2.11 | 0.098 |
| *LHA* | HY | 2.08 | 0.055 |
| *PAG* | MB | 1.89 | 0.067 |

**Table S3.**
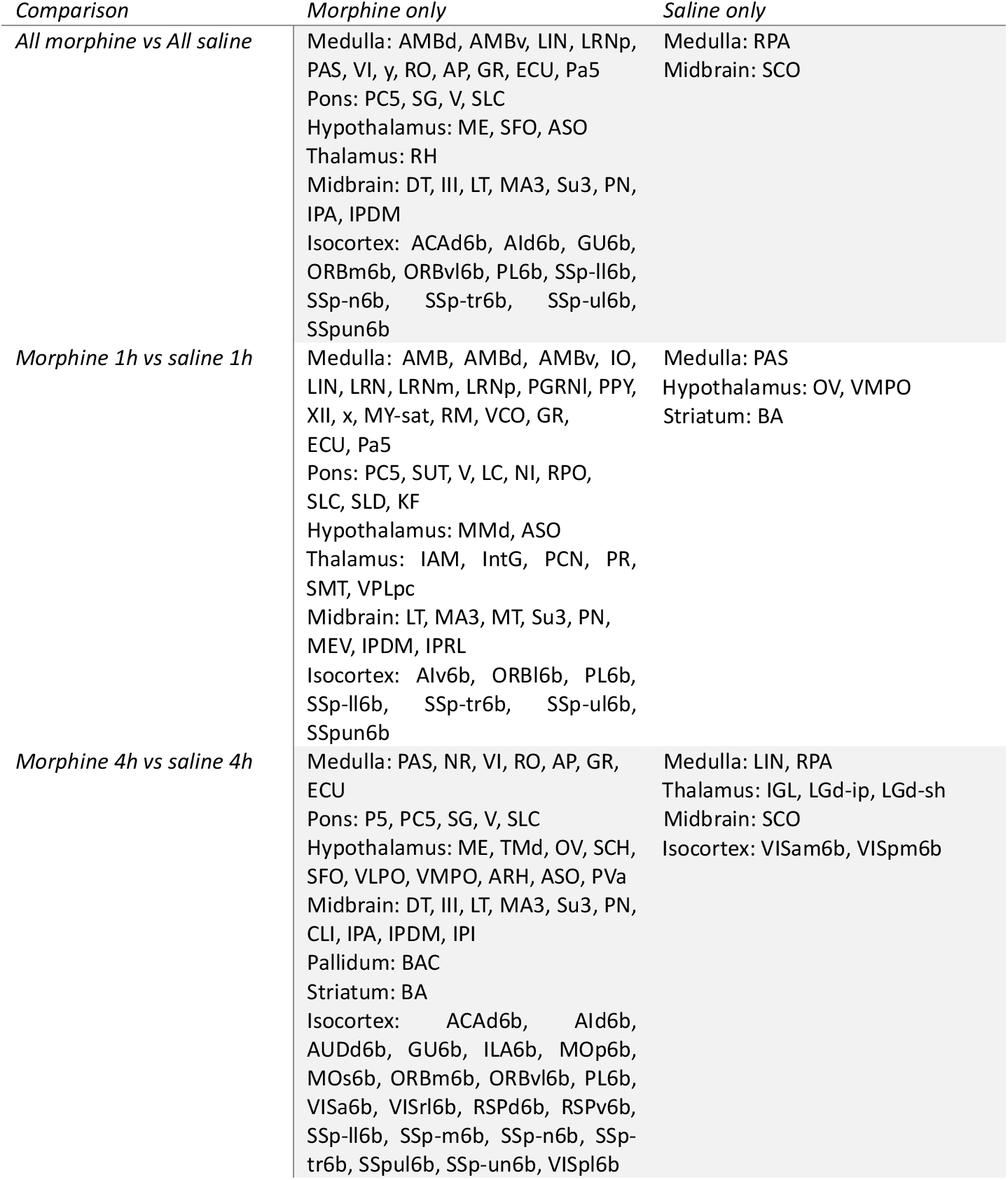
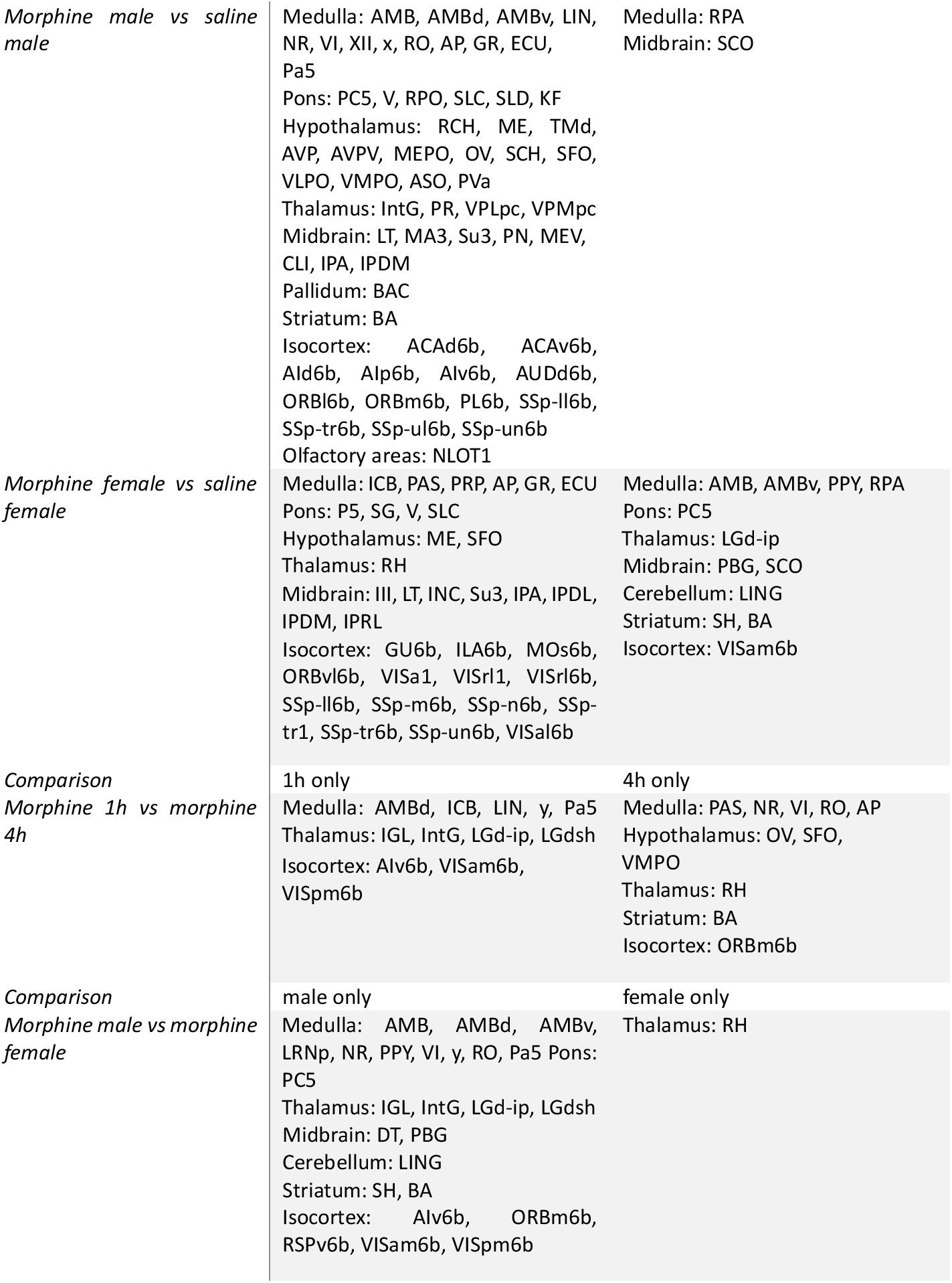
Binary Effects: Regions of c-Fos activity in the brain which are only present in one experimental group.

**Table S4.** Structures with significant differences in c-Fos+ cell densities (morphine vs saline) at 1h (sorted by decreasing density fold-change) *ICc* 2.23 0.0306

| *Acronym* | *Density*<br>*Ratio* | *p-value* |
| --- | --- | --- |
| *PB* | 11.11 | 0.0444 |
| *EPd* | 5.27 | 0.0235 |
| *ENTl5* | 4.37 | 0.008 |
| *ENTm3* | 3.94 | 0.0409 |
| *ENTm5* | 3.81 | 0.0347 |
| *VISpor5* | 3.51 | 0.0413 |
| *AUDd5* | 3.48 | 0.0197 |
| *SSs5* | 3.42 | 0.0365 |
| *VISp5* | 3.40 | 0.031 |
| *ENTm6* | 3.14 | 0.0374 |
| *MRN* | 3.08 | 0.0462 |
| *CA3* | 2.61 | 0.0281 |

**Table S5.**
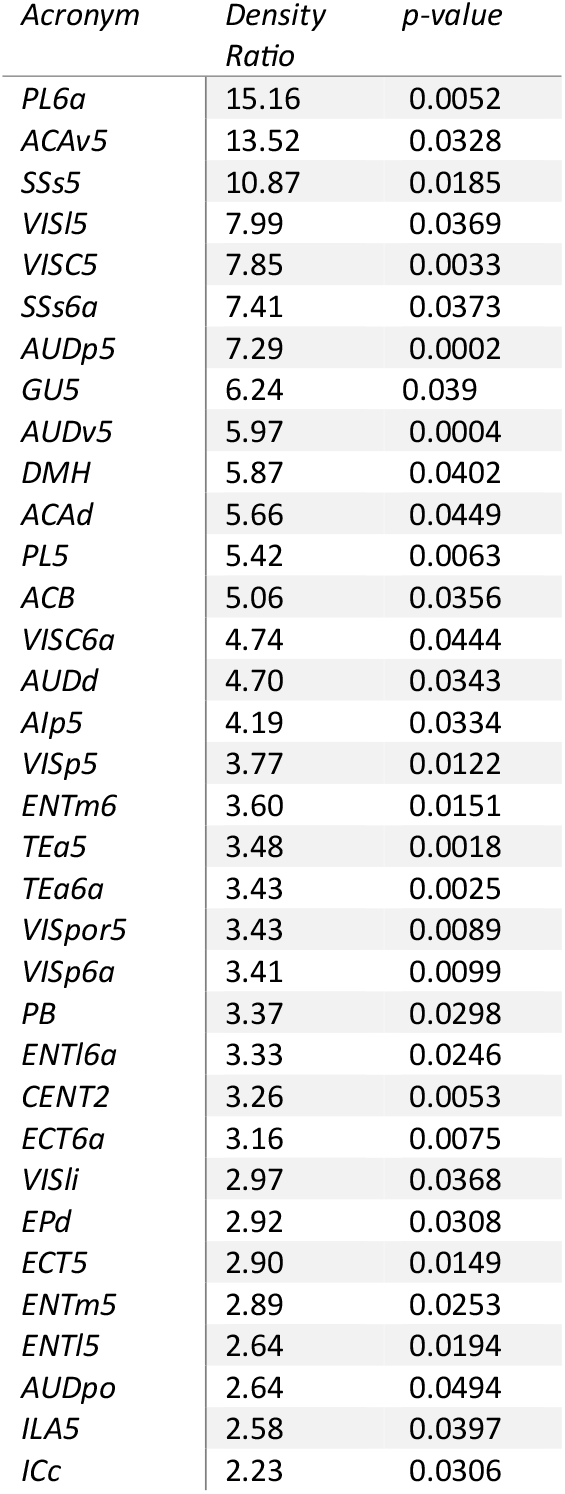
Structures with significant differences in c-Fos+ cell densities (morphine vs saline) at 4h (sorted by decreasing fold-change)

**Table S6.** Male-specific morphine effects. Structures with significant differences in c-Fos+ cell densities (morphine/saline) in male mice (sorted by decreasing fold-change)

| *Acronym* | *Density Ratio* | *p-value* |
| --- | --- | --- |
| *ACB* | 15.6 | 0.0098 |
| *ACAv5* | 10.29 | 0.0486 |
| *LA* | 9.47 | 0.0247 |
| *PAL* | 8.49 | 0.035 |
| *EPd* | 7.83 | 0.0089 |
| *BLAp* | 7.72 | 0.0176 |
| *AUDd5* | 7 | 0.0064 |
| *PL6a* | 6.91 | 0.0295 |
| *ACAd6a* | 6.86 | 0.045 |
| *VISI5* | 5.95 | 0.0273 |
| *ECT6a* | 5.82 | 0.0141 |
| *PL5* | 5.77 | 0.0363 |
| *SSs5* | 5.52 | 0.0054 |
| *BMAp* | 5.4 | 0.0255 |
| *ENTI5* | 5.15 | 0.0121 |
| *LHA* | 5.14 | 0.0314 |
| *PB* | 5.04 | 0.0303 |
| *TEa6a* | 5 | 0.05 |
| *ENTm5* | 4.86 | 0.0153 |
| *ENTm6* | 4.83 | 0.0098 |
| *Alp5* | 4.72 | 0.027 |
| *PA* | 4.54 | 0.0156 |
| *VISC5* | 4.36 | 0.0412 |
| *ENTl6a* | 4.25 | 0.0256 |
| *VISpor5* | 3.48 | 0.0491 |
| *AUDp5* | 3.14 | 0.0385 |
| *AUDv5* | 3.14 | 0.0322 |
| *CENT2* | 2.96 | 0.0413 |
| *CA3* | 2.53 | 0.0341 |
| *ProS* | 2.32 | 0.0162 |
| *CA1* | 2.15 | 0.0399 |
| *SUB* | 1.91 | 0.0431 |

**Table S7.**
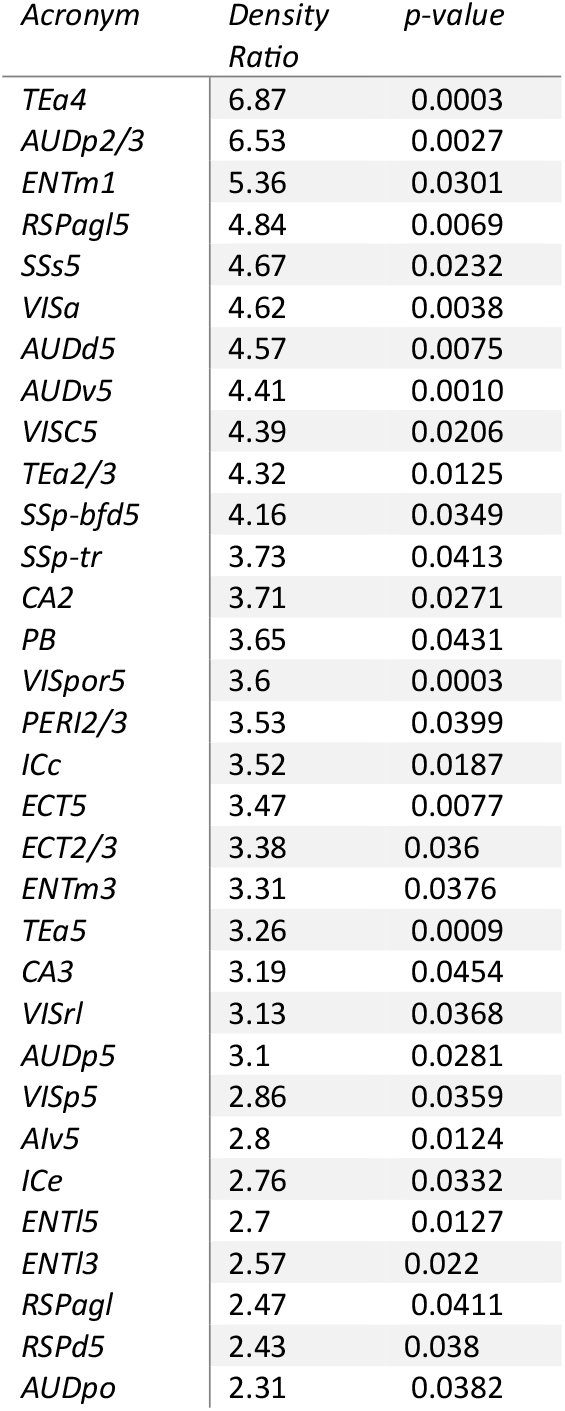
Female-specific morphine effects. Structures with significant differences in c-Fos+ cell densities (morphine/saline) in female mice (sorted by decreasing fold-change)

| *Acronym* | *Density*<br>*Ratio* | *p-value* |
| --- | --- | --- |
| *TEa4* | 6.87 | 0.0003 |
| *AUDp2/3* | 6.53 | 0.0027 |
| *ENTm1* | 5.36 | 0.0301 |
| *RSPagl5* | 4.84 | 0.0069 |
| *SSs5* | 4.67 | 0.0232 |
| *VISa* | 4.62 | 0.0038 |
| *AUDd5* | 4.57 | 0.0075 |
| *AUDv5* | 4.41 | 0.0010 |
| *VISC5* | 4.39 | 0.0206 |
| *TEa2/3* | 4.32 | 0.0125 |
| *SSp-bfd5* | 4.16 | 0.0349 |
| *SSp-tr* | 3.73 | 0.0413 |
| *CA2* | 3.71 | 0.0271 |
| *PB* | 3.65 | 0.0431 |
| *VISpor5* | 3.6 | 0.0003 |
| *PERl2/3* | 3.53 | 0.0399 |
| *ICc* | 3.52 | 0.0187 |
| *ECT5* | 3.47 | 0.0077 |
| *ECT2/3* | 3.38 | 0.036 |
| *ENTm3* | 3.31 | 0.0376 |
| *TEa5* | 3.26 | 0.0009 |
| *CA3* | 3.19 | 0.0454 |
| *VISrl* | 3.13 | 0.0368 |
| *AUDp5* | 3.1 | 0.0281 |
| *VISp5* | 2.86 | 0.0359 |
| *Alv5* | 2.8 | 0.0124 |
| *ICe* | 2.76 | 0.0332 |
| *ENTl5* | 2.7 | 0.0127 |
| *ENTl3* | 2.57 | 0.022 |
| *RSPagl* | 2.47 | 0.0411 |
| *RSPd5* | 2.43 | 0.038 |
| *AUDpo* | 2.31 | 0.0382 |

**Table S8.**
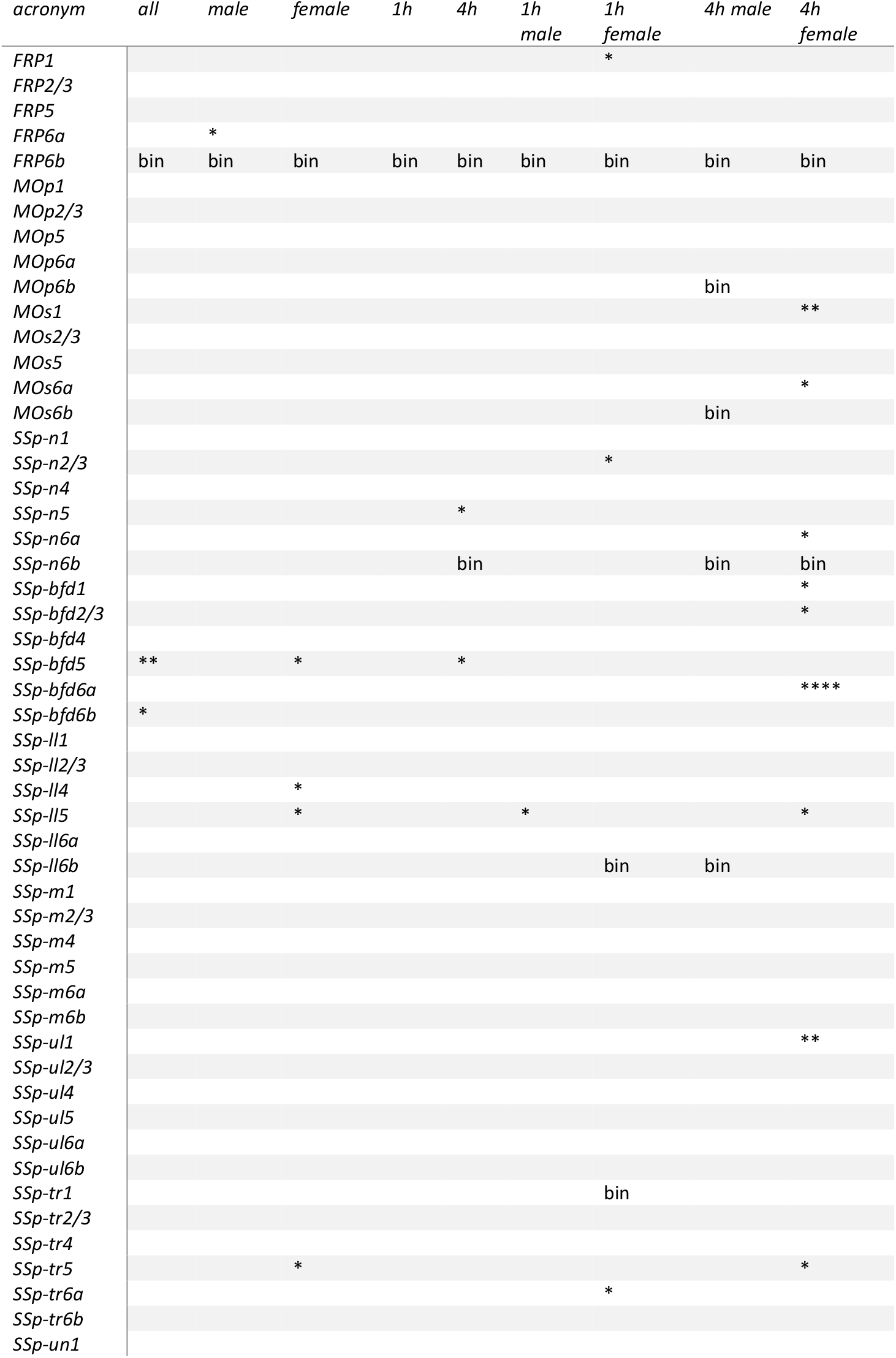
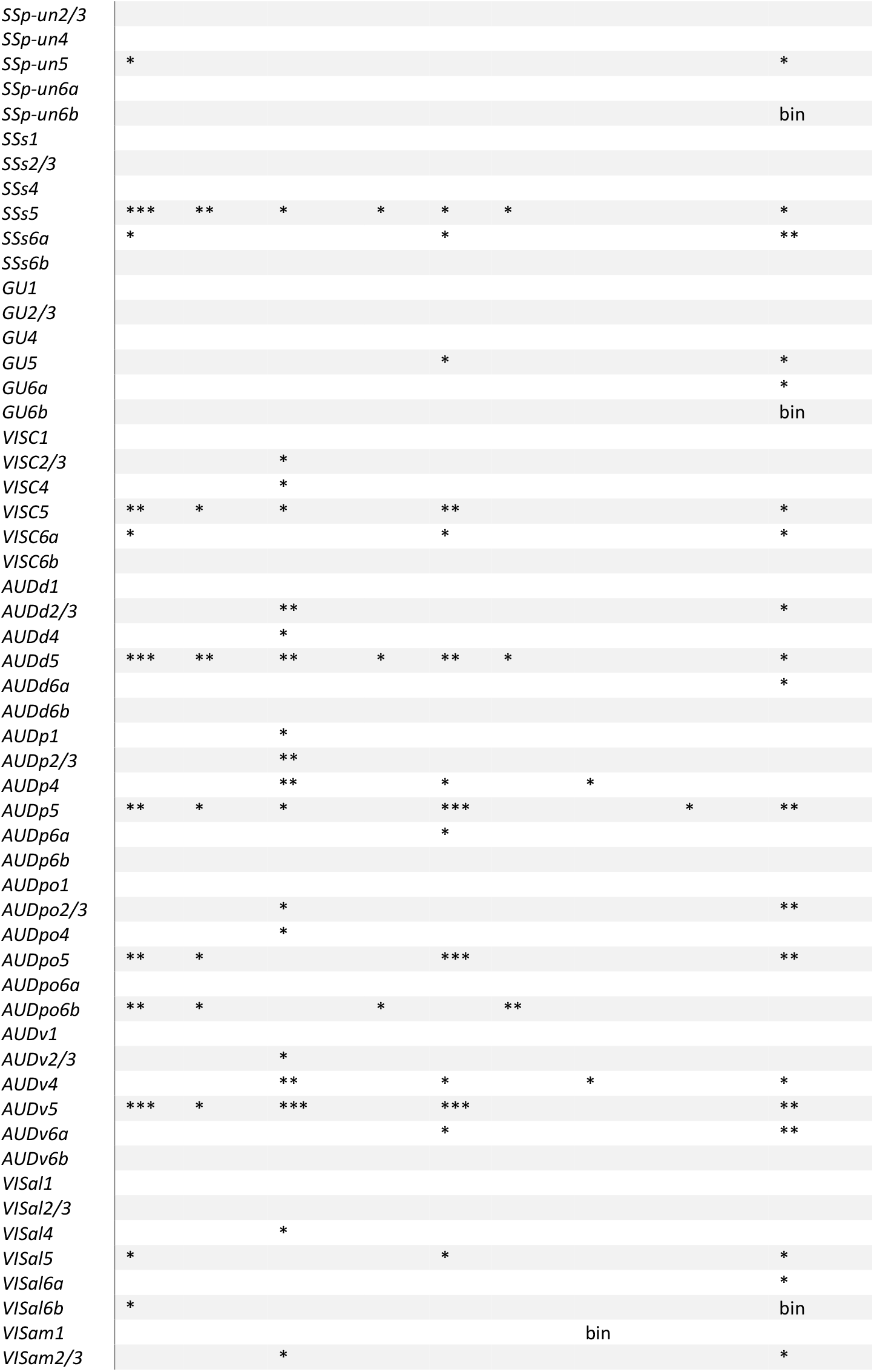
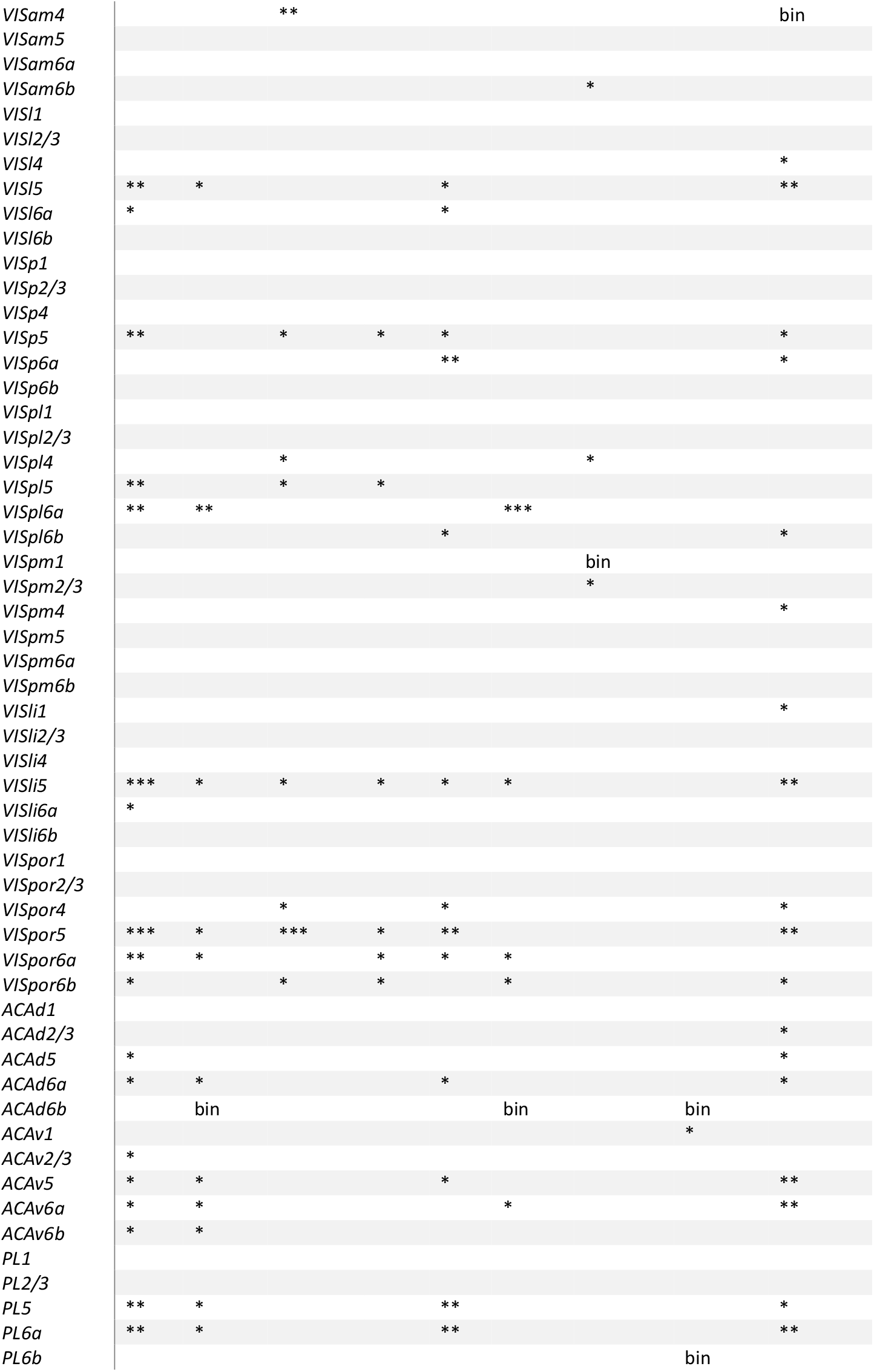
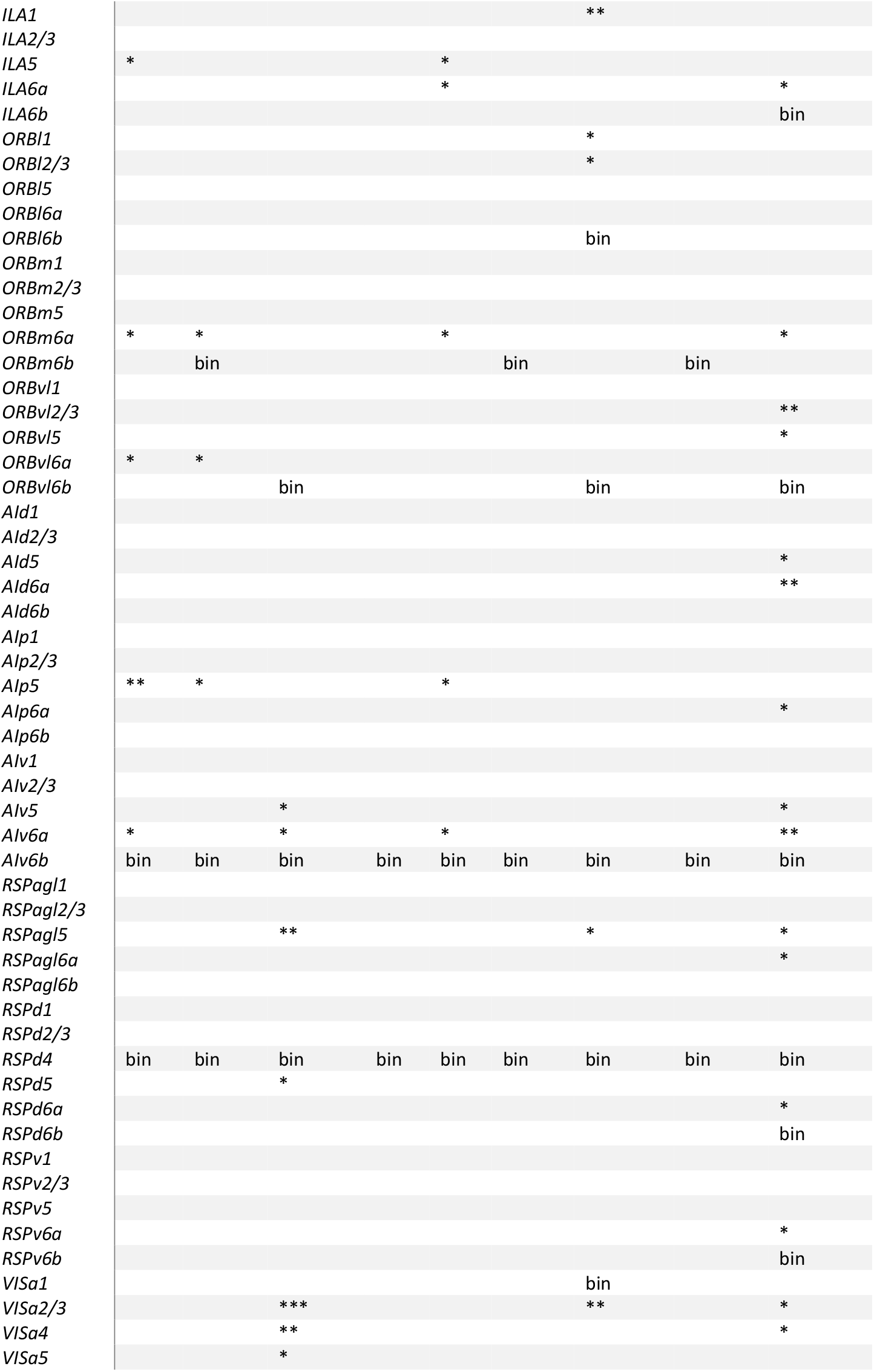
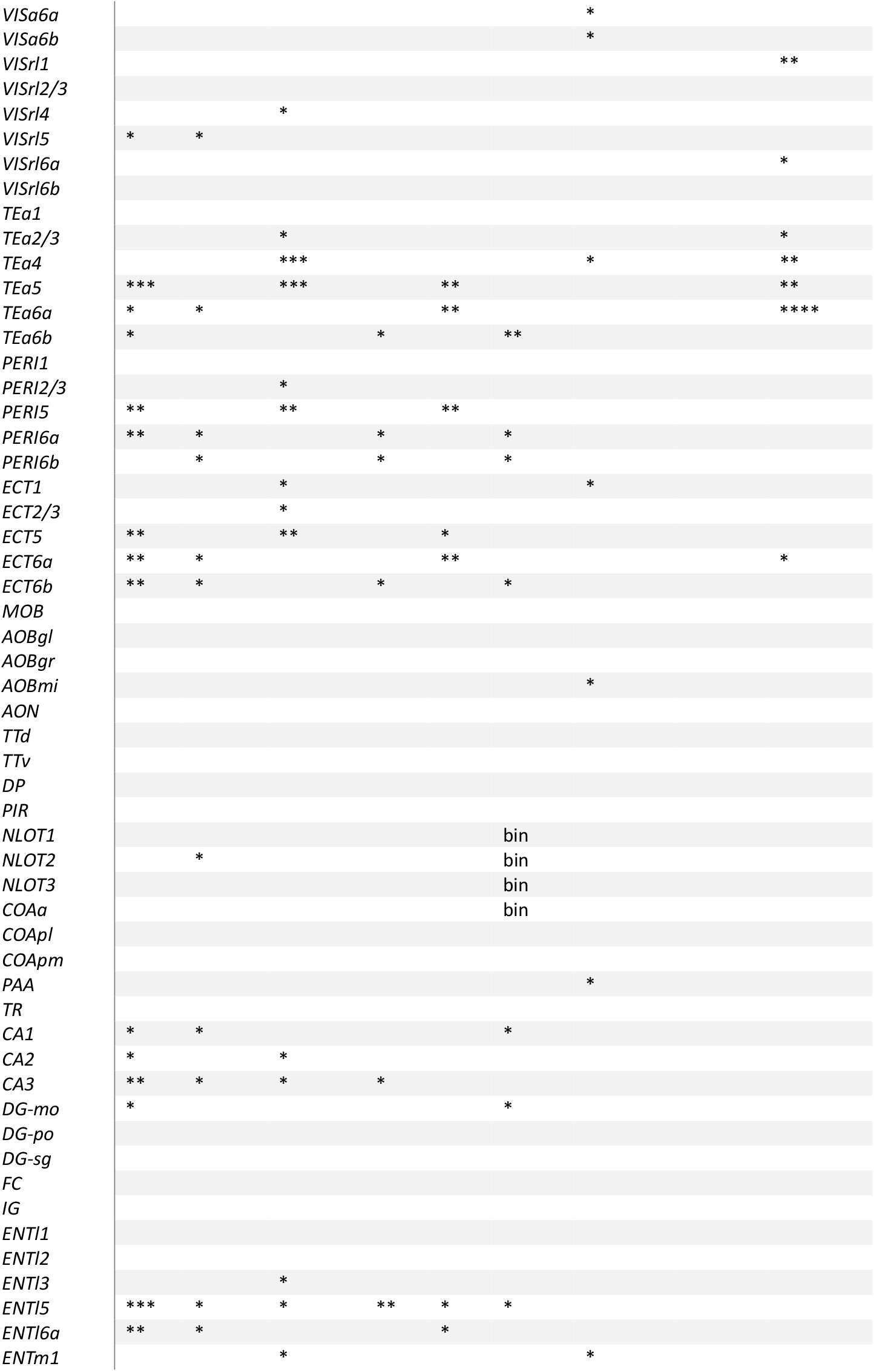
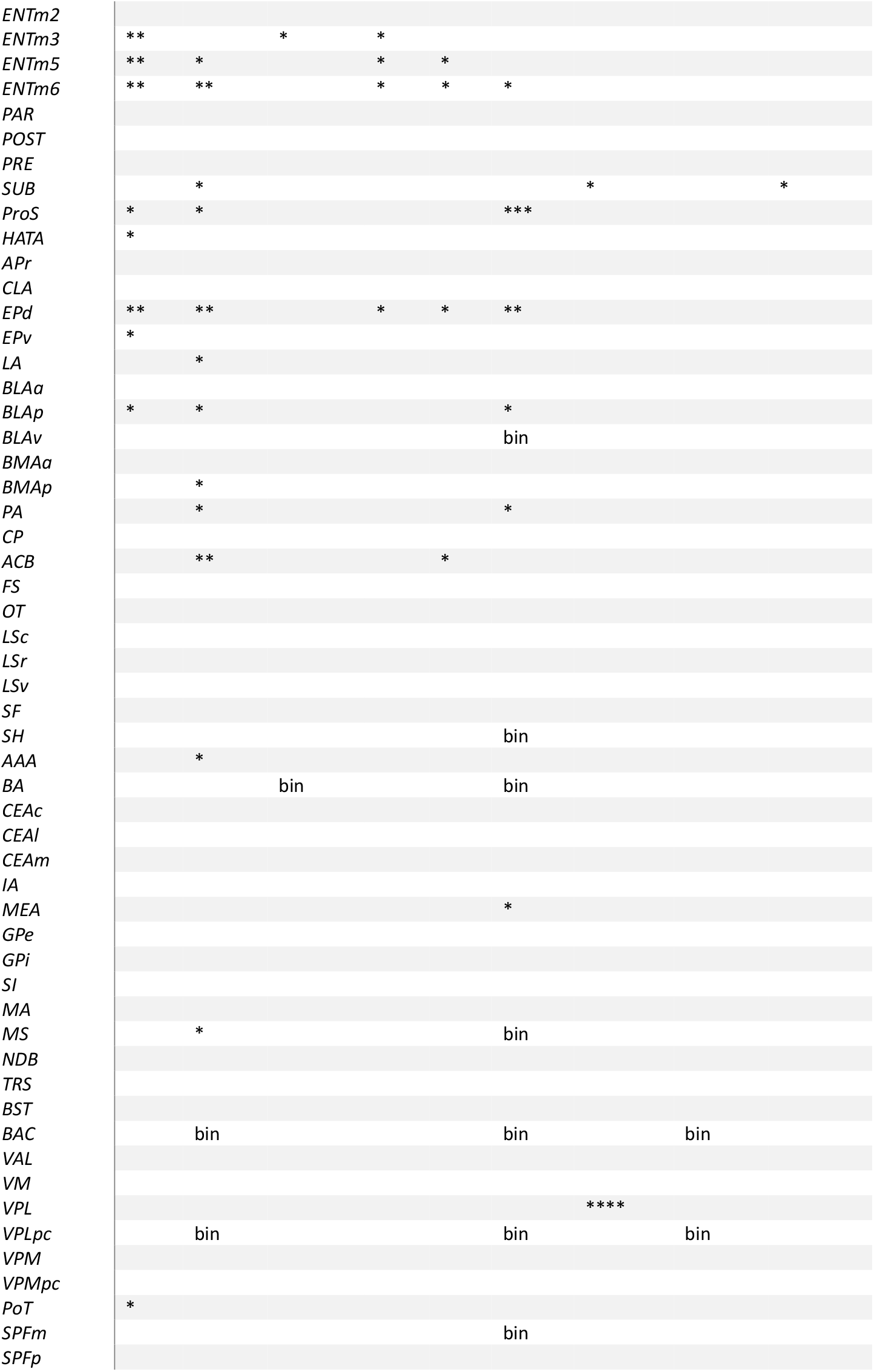
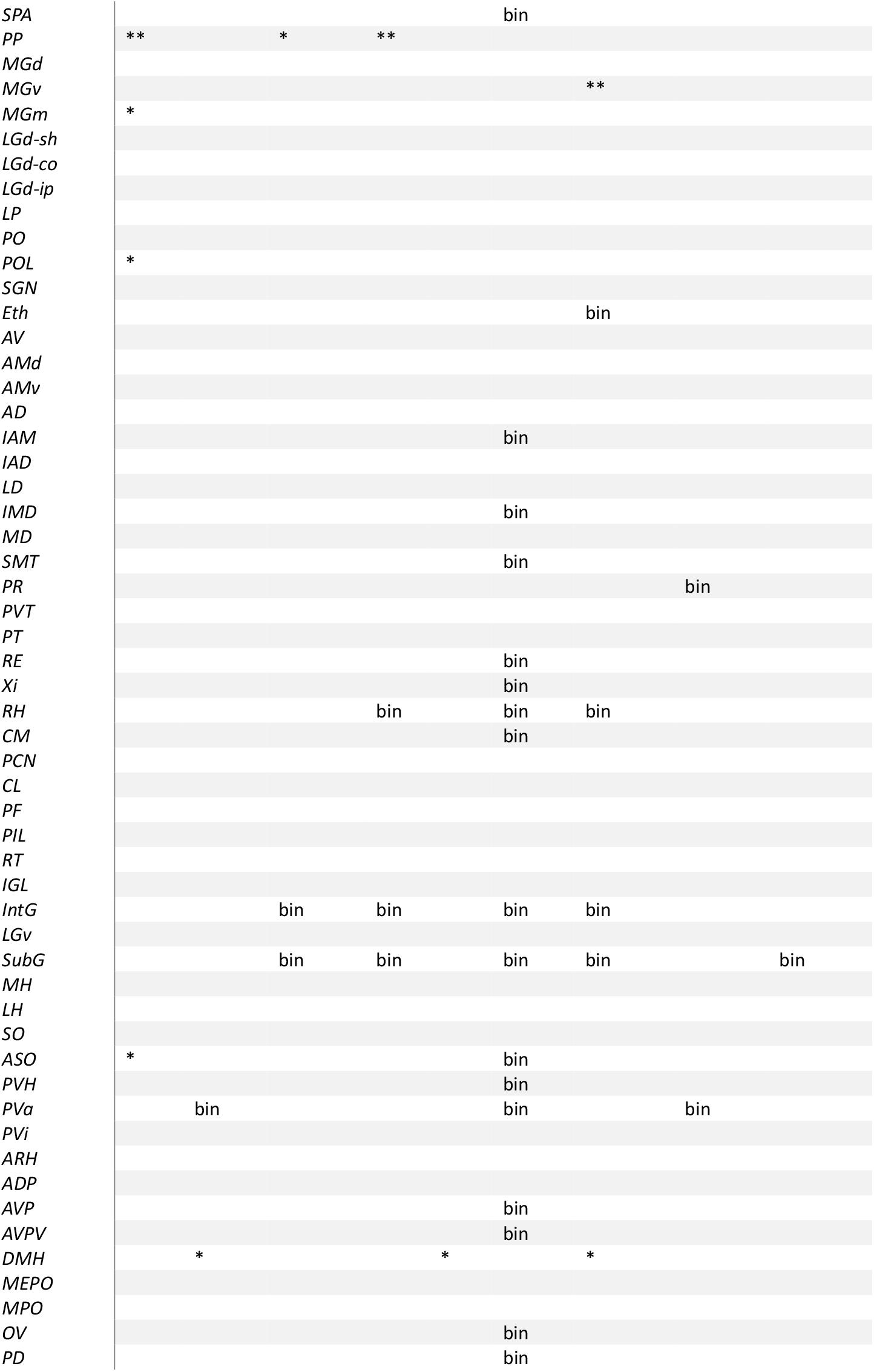
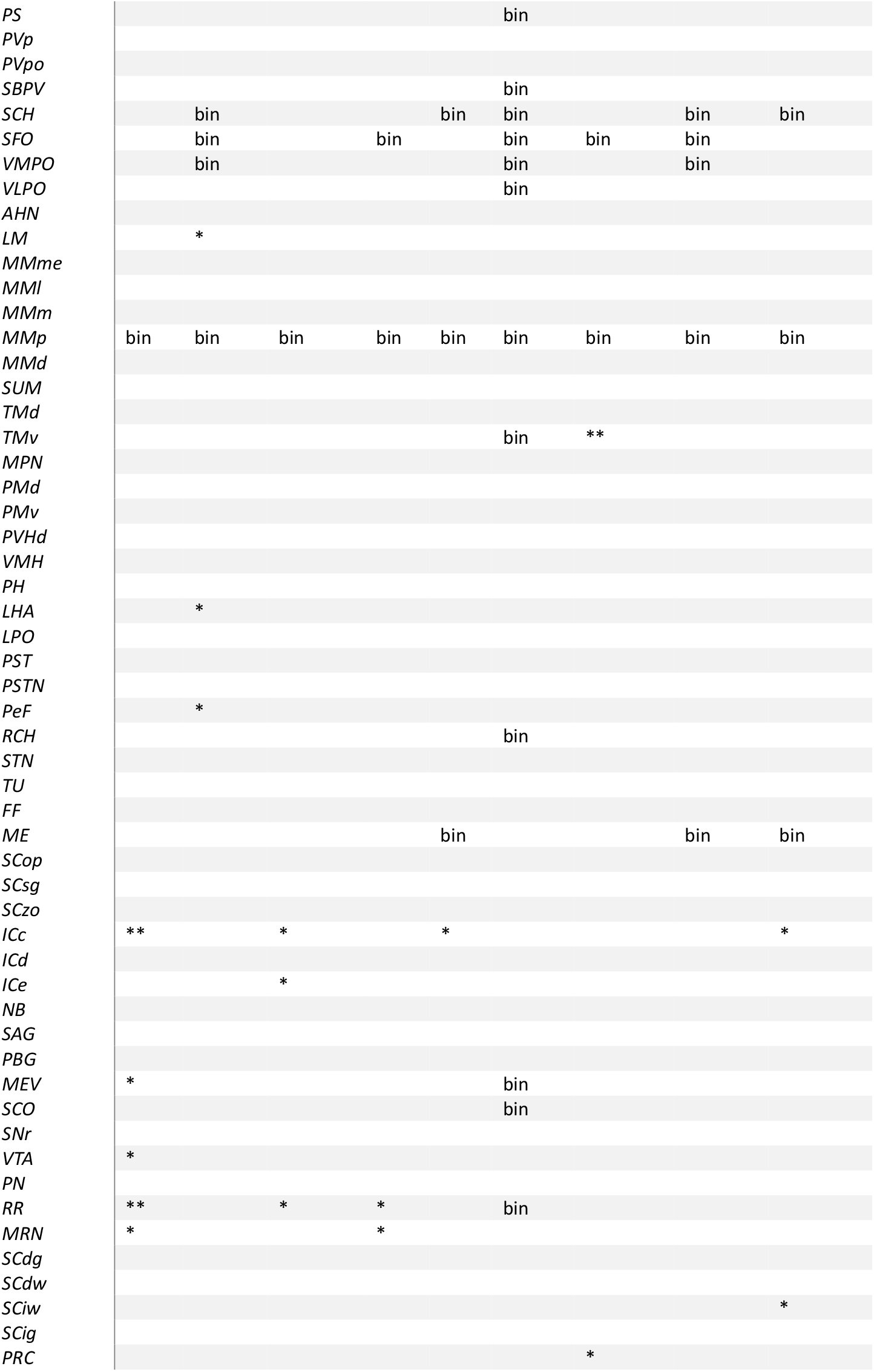
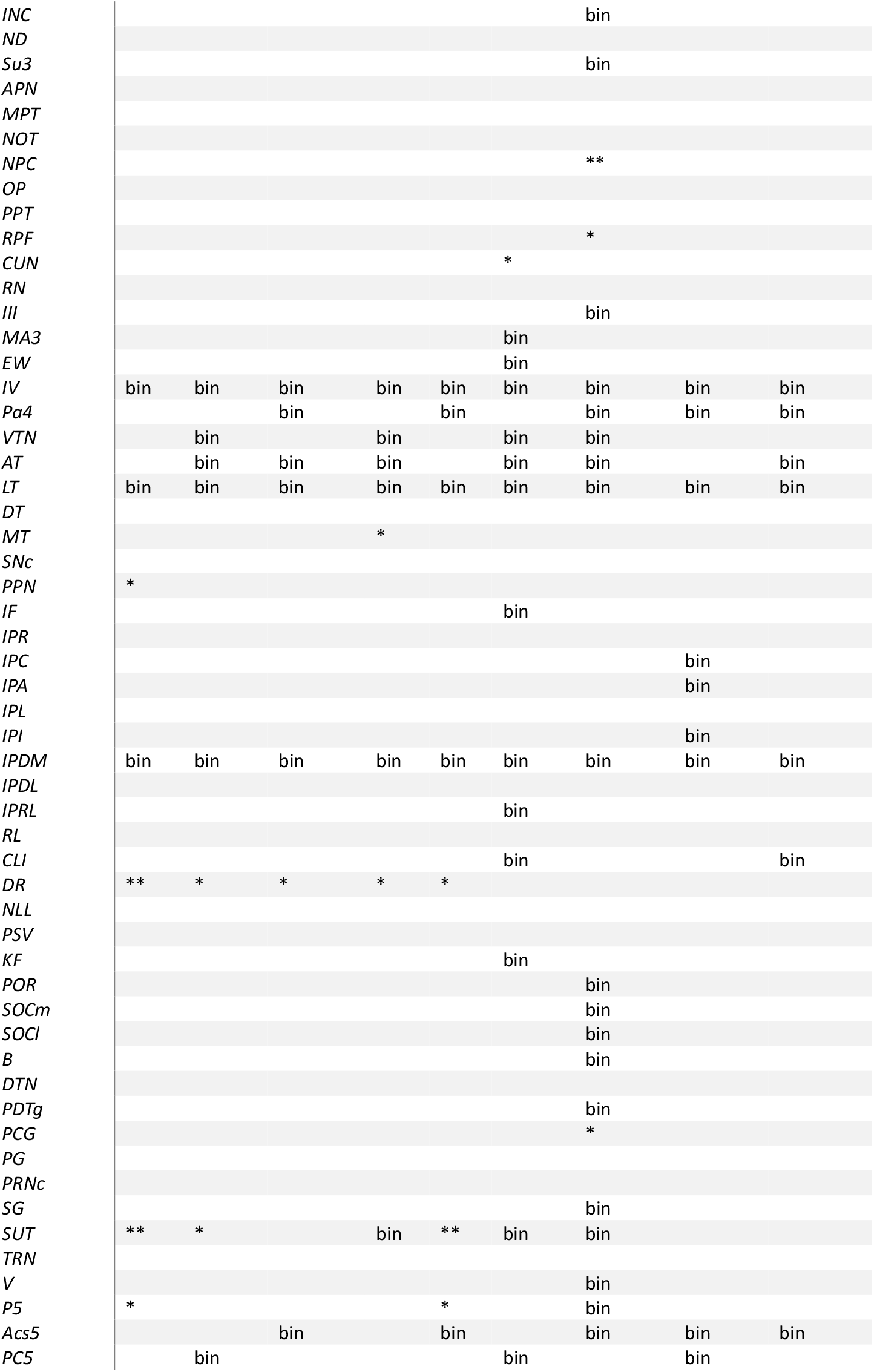
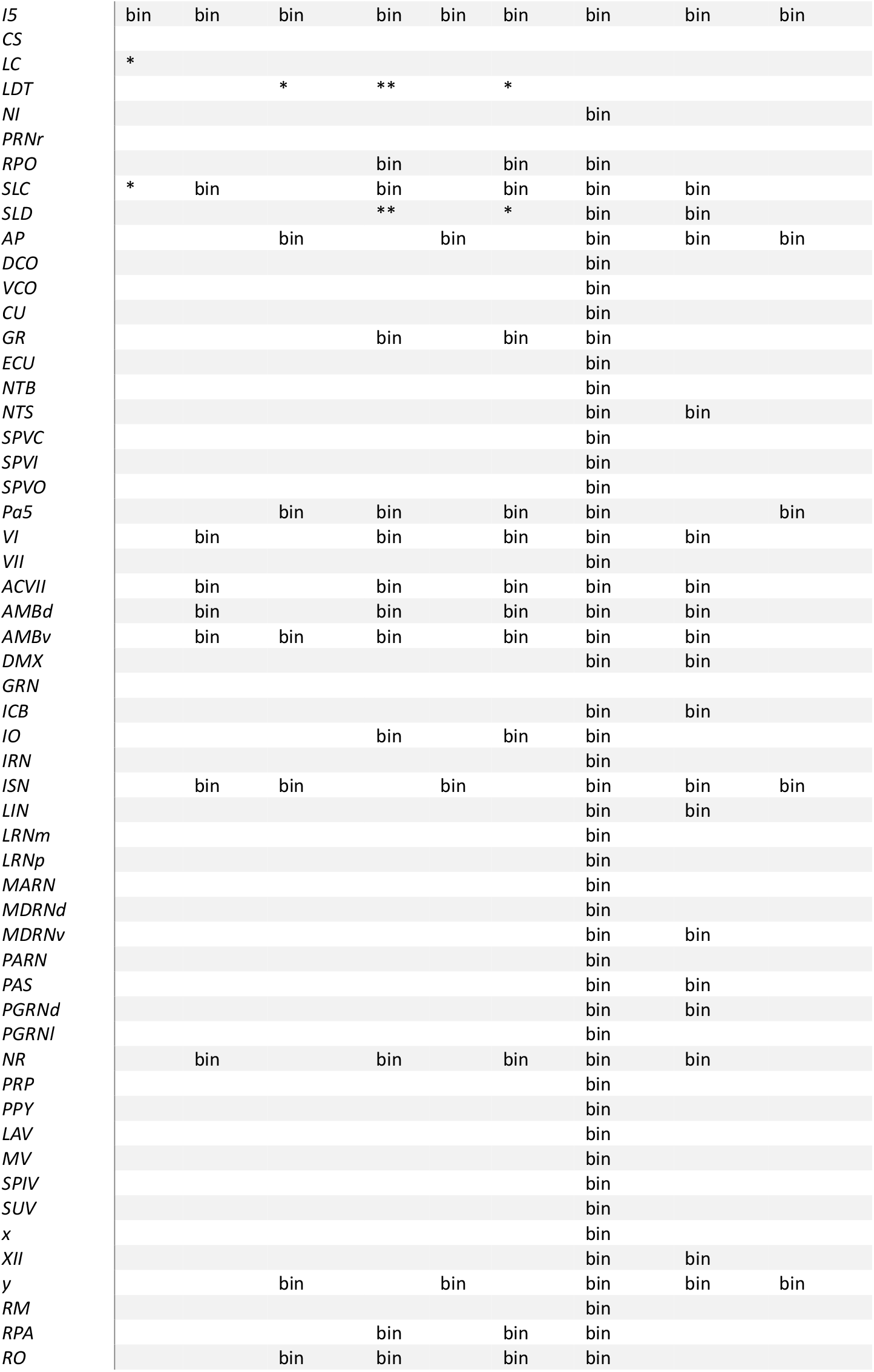
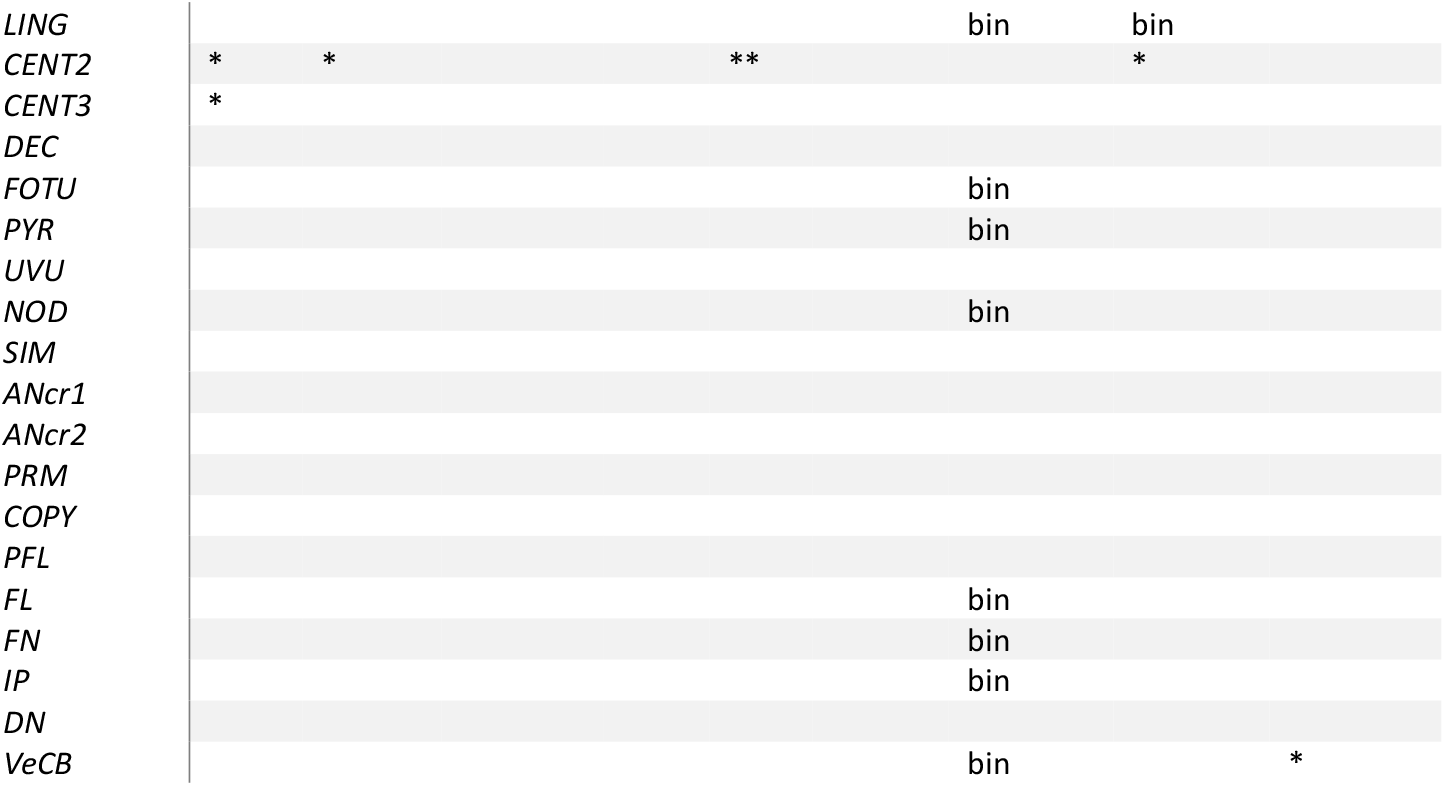
Significant effects of morphine on all leaf structures within the CCFv3 tree containing grey matter. In each column, significance was calculated between respective morphine and saline subgroups.

**Figure S3.**
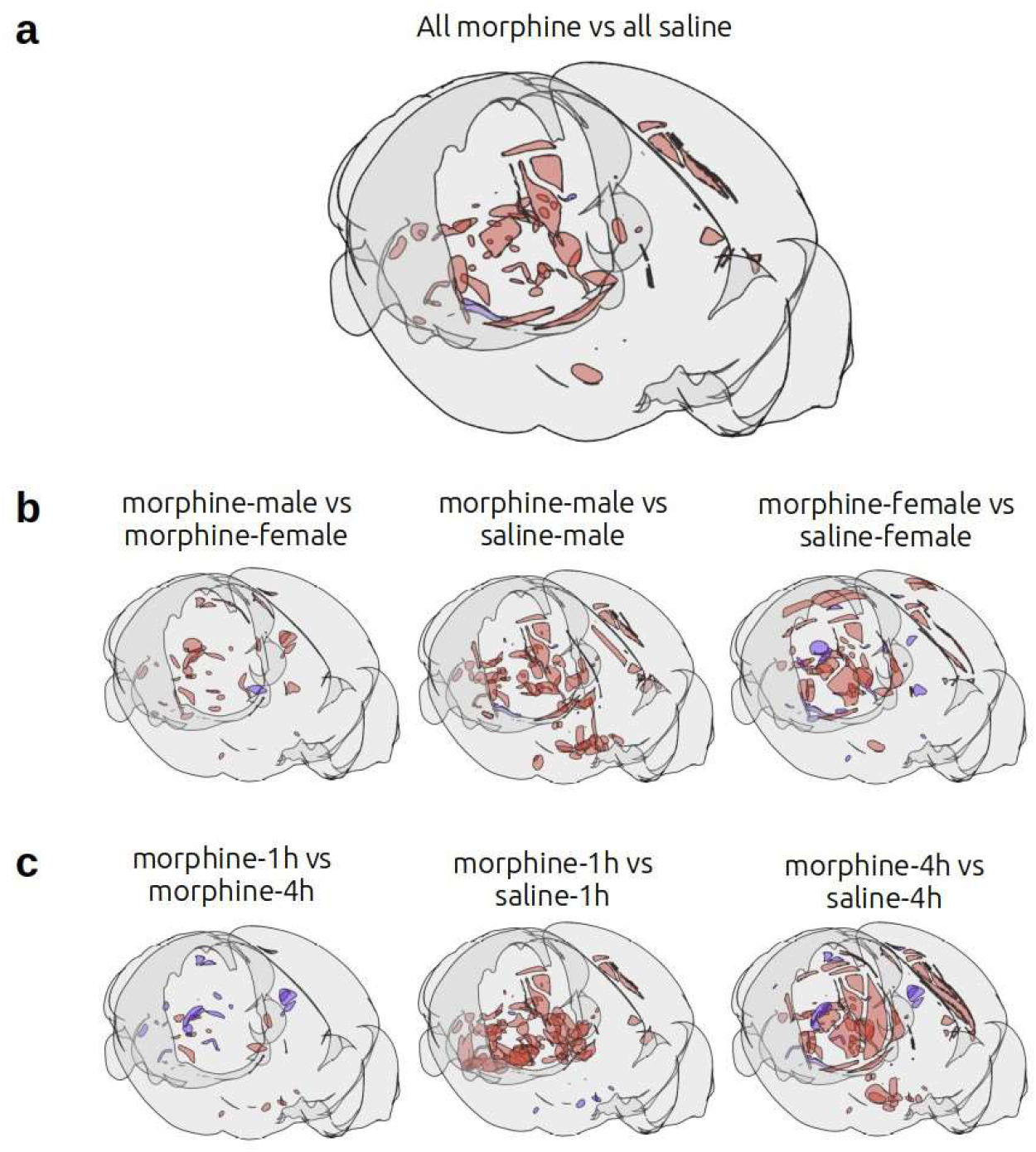
Binary Effects. Cartoon visualization of brain regions with c-Fos activity in the brain which are only present in one experimental group. **a)** red = morphine only, blue = saline only; **b) left:** red = male only, blue = female only; **middle, right:** red = morphine only, blue = saline only; **c) left:** red = 1h only, blue = 4h only; **middle, right:** red = morphine only, blue = saline only.

**Figure S4.**
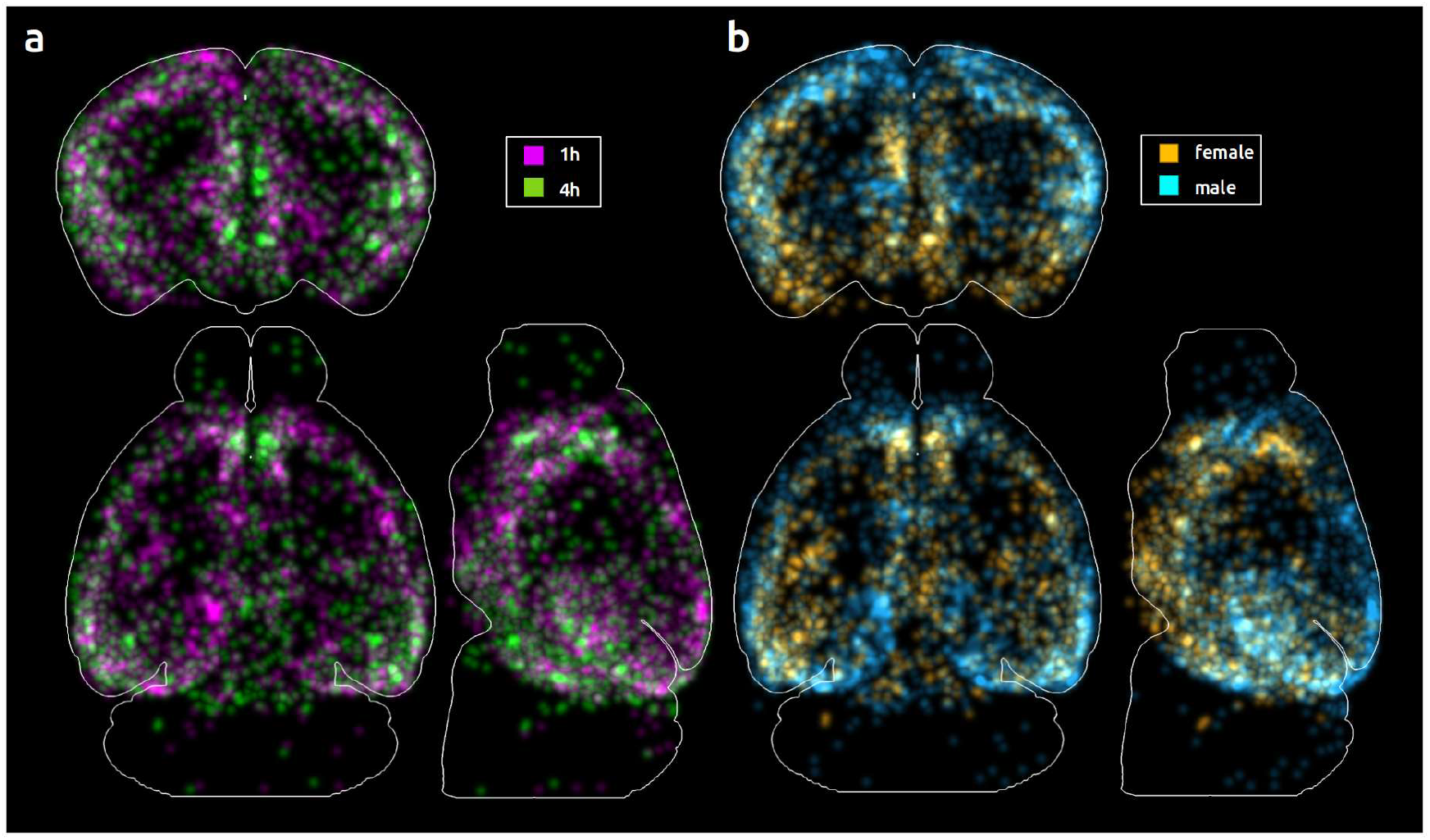
Temporally and sexually dimorphic activation patterns after morphine exposure: Maximum intensity projections of c-Fos puncta locations in the CCFv3 for morphine brains normalized (divided) by corresponding saline controls (smoothed with Gaussian filter, kernel size=10)

## References

1. Koob, G. F. Neurobiology of Opioid Addiction: Opponent Process, Hyperkatifeia, and Negative Reinforcement. Biol Psychiatry 87, 44–53 (2020).

2. Jones, A. A., Segel, J. E., Skogseth, E. M., Apsley, H. B. & Santos-Lozada, A. R. Drug overdose deaths among women 1999-2021 in the United States: Differences by race, ethnicity, and age. Womens Health (Lond) 20, 17455057241307088 (2024).

3. Tiwari, K., Rahimian, M. A., Roberts, M. S., Kumar, P. & Buchanich, J. M. Measuring network dynamics of opioid overdose deaths in the United States. Sci Rep 14, 29563 (2024).

4. Jalal, H. et al. Changing dynamics of the drug overdose epidemic in the United States from 1979 through 2016. Science 361, eaau1184 (2018).

5. Gossop, M., Green, L., Phillips, G. & Bradley, B. Lapse, relapse and survival among opiate addicts after treatment. A prospective follow-up study. Br J Psychiatry 154, 348–353 (1989).

6. Harris, A. C. & Gewirtz, J. C. Acute opioid dependence: characterizing the early adaptations underlying drug withdrawal. Psychopharmacology (Berl) 178, 353–366 (2005).

7. Cruz, F. C., Javier Rubio, F. & Hope, B. T. Using c-fos to study neuronal ensembles in corticostriatal circuitry of addiction. Brain Res 1628, 157–173 (2015).

8. Cruz, F. C. et al. Role of nucleus accumbens shell neuronal ensembles in context-induced reinstatement of cocaine-seeking. J Neurosci 34, 7437–7446 (2014).

9. Whitaker, L. R. & Hope, B. T. Chasing the addicted engram: identifying functional alterations in Fos-expressing neuronal ensembles that mediate drug-related learned behavior. Learn Mem 25, 455–460 (2018).

10. Salery, M., Godino, A. & Nestler, E. J. Drug-activated cells: From immediate early genes to neuronal ensembles in addiction. Adv Pharmacol 90, 173–216 (2021).

11. Koya, E., Margetts-Smith, G. & Hope, B. T. Daun02 Inactivation of Behaviorally Activated Fos-Expressing Neuronal Ensembles. Curr Protoc Neurosci 76, 8.36.1-8.36.17 (2016).

12. Warren, B. L., Suto, N. & Hope, B. T. Mechanistic Resolution Required to Mediate Operant Learned Behaviors: Insights from Neuronal Ensemble-Specific Inactivation. Front Neural Circuits 11, 28 (2017).

13. Muntifering, M. et al. Clearing for Deep Tissue Imaging. Curr Protoc Cytom 86, e38 (2018).

14. Watson, A. M. et al. Ribbon scanning confocal for high-speed high-resolution volume imaging of brain. PLoS One 12, e0180486 (2017).

15. FAIR Principles. GO FAIR https://www.go-fair.org/fair-principles/.

16. Wilkinson, M. D. et al. The FAIR Guiding Principles for scientific data management and stewardship. Sci Data 3, 160018 (2016).

17. Wang, Q. et al. The Allen Mouse Brain Common Coordinate Framework: A 3D Reference Atlas. Cell 181, 936-953.e20 (2020).

18. Yoo, A. B., Jette, M. A. & Grondona, M. SLURM: Simple Linux Utility for Resource Management. In Job Scheduling Strategies for Parallel Processing (eds. Feitelson, D., Rudolph, L. & Schwiegelshohn, U.) vol. 2862 44–60 (Springer Berlin Heidelberg, Berlin, Heidelberg, 2003).

19. Jacquet, Y. F. & Lajtha, A. The periaqueductal gray: site of morphine analgesia and tolerance as shown by 2-way cross tolerance between systemic and intracerebral injections. Brain Res 103, 501–513 (1976).

20. Ikemoto, S. et al. Brain reward circuitry beyond the mesolimbic dopamine system: a neurobiological theory. Neurosci Biobehav Rev 35, 129–150 (2010).

21. Welsch, L. et al. Mu Opioid Receptor-Expressing Neurons in the Dorsal Raphe Nucleus Are Involved in Reward Processing and Affective Behaviors. Biol Psychiatry 94, 842– 851 (2023).

22. Miyachi, S. et al. [Cortico-basal ganglia circuits--parallel closed loops and convergent/divergent connections]. Brain Nerve 61, 351–359 (2009).

23. Haber, S. N. & Calzavara, R. The cortico-basal ganglia integrative network: the role of the thalamus. Brain Res Bull 78, 69–74 (2009).

24. Haber, S. N. & Knutson, B. The reward circuit: linking primate anatomy and human imaging. Neuropsychopharmacology 35, 4–26 (2010).

25. Kibaly, C., Xu, C., Cahill, C. M., Evans, C. J. & Law, P.-Y. et al. Nonnociceptive roles of opioids in the CNS: opioids’ effects on neurogenesis, learning, memory and affect. Nat Rev Neurosci 20, 5–18 (2019).

26. Fredriksson, I. et al. Role of ventral subiculum neuronal ensembles in incubation of oxycodone craving after electric barrier-induced voluntary abstinence. Sci Adv 9, eadd8687 (2023).

27. Ceceli, A. O., Bradberry, C. W. & Goldstein, R. Z. The neurobiology of drug addiction: cross-species insights into the dysfunction and recovery of the prefrontal cortex. Neuropsychopharmacology 47, 276–291 (2022).

28. McGregor, M. S. & LaLumiere, R. T. Still a ‘hidden island’? The rodent insular cortex in drug seeking, reward, and risk. Neurosci Biobehav Rev 153, 105334 (2023).

29. McGregor, M. S., Cosme, C. V. & LaLumiere, R. T. Insular cortex subregions have distinct roles in cued heroin seeking after extinction learning and prolonged withdrawal in rats. Neuropsychopharmacol. 49, 1540–1549 (2024).

30. McKendrick, G., McDevitt, D. S., Shafeek, P., Cottrill, A. & Graziane, N. M. Anterior cingulate cortex and its projections to the ventral tegmental area regulate opioid withdrawal, the formation of opioid context associations and context-induced drug seeking. Front Neurosci 16, 972658 (2022).

31. Gf, K. & Nd, V. Neurocircuitry of addiction. Neuropsychopharmacology : official publication of the American College of Neuropsychopharmacology 35, (2010).

32. Sheng, H. et al. Nucleus accumbens circuit disinhibits lateral hypothalamus glutamatergic neurons contributing to morphine withdrawal memory in male mice. Nat Commun 14, 71 (2023).

33. Baladron, J. & Hamker, F. H. Habit learning in hierarchical cortex-basal ganglia loops. Eur J Neurosci 52, 4613–4638 (2020).

34. Salamone, J. D. & Correa, M. The mysterious motivational functions of mesolimbic dopamine. Neuron 76, 470–485 (2012).

35. Kaplan, G. B. & Thompson, B. L. Neuroplasticity of the extended amygdala in opioid withdrawal and prolonged opioid abstinence. Front Pharmacol 14, 1253736 (2023).

36. Ammon-Treiber, S. & Höllt, V. et al. Morphine-induced changes of gene expression in the brain. Addict Biol 10, 81–89 (2005).

37. E, E. et al. A mu-delta opioid receptor brain atlas reveals neuronal co-occurrence in subcortical networks. Brain structure & function 220, (2015).

38. Fatt, M. P. et al. Morphine-responsive neurons that regulate mechanical antinociception. Science 385, eado6593 (2024).

39. Tao, R. & Auerbach, S. B. Involvement of the dorsal raphe but not median raphe nucleus in morphine-induced increases in serotonin release in the rat forebrain. Neuroscience 68, 553–561 (1995).

40. Tao, R. & Auerbach, S. B. GABAergic and glutamatergic afferents in the dorsal raphe nucleus mediate morphine-induced increases in serotonin efflux in the rat central nervous system. J Pharmacol Exp Ther 303, 704–710 (2002).

41. Proudfit, H. K. et al. Effects of raphe magnus and raphe pallidus lesions on morphine-induced analgesia and spinal cord monoamines. Pharmacol Biochem Behav 13, 705–714 (1980).

42. Kalyuzhny, A. E., Arvidsson, U., Wu, W. & Wessendorf, M. W. mu-Opioid and delta-opioid receptors are expressed in brainstem antinociceptive circuits: studies using immunocytochemistry and retrograde tract-tracing. J Neurosci 16, 6490–6503 (1996).

43. Huhn, A. S., Berry, M. S. & Dunn, K. E. Systematic review of sex-based differences in opioid-based effects. Int Rev Psychiatry 30, 107–116 (2018).

44. Sharp, J. L., Pearson, T. & Smith, M. A. Sex differences in opioid receptor mediated effects: Role of androgens. Neurosci Biobehav Rev 134, 104522 (2022).

45. Wj, L. & Me, C. Sex differences in the acquisition of intravenously self-administered cocaine and heroin in rats. Psychopharmacology 144, (1999).

46. Carroll, M. E., Morgan, A. D., Lynch, W. J., Campbell, U. C. & Dess, N. K. Intravenous cocaine and heroin self-administration in rats selectively bred for differential saccharin intake: phenotype and sex differences. Psychopharmacology (Berl) 161, 304–313 (2002).

47. Cicero, T. J., Aylward, S. C. & Meyer, E. R. Gender differences in the intravenous self-administration of mu opiate agonists. Pharmacol Biochem Behav 74, 541–549 (2003).

48. Klein, L. C., Popke, E. J. & Grunberg, N. E. Sex differences in effects of predictable and unpredictable footshock on fentanyl self-administration in rats. Exp Clin Psychopharmacol 5, 99–106 (1997).

49. Lee, C.W.-S. & Ho, I.-K. et al. Sex differences in opioid analgesia and addiction: interactions among opioid receptors and estrogen receptors. Mol Pain 9, 45 (2013).

50. Wu, L.-T. & Howard, M. O. Is inhalant use a risk factor for heroin and injection drug use among adolescents in the United States? Addict Behav 32, 265–281 (2007).

51. M, Z. et al. Distinct and sex-specific expression of mu opioid receptors in anterior cingulate and somatosensory S1 cortical areas. Pain 164, (2023).

52. Sb, E. & Ma, S. Estradiol and Mu opioid-mediated reward: The role of estrogen receptors in opioid use. Addiction neuroscience 9, (2023).

53. Hodgson, S. R., Hofford, R. S., Roberts, K. W., Wellman, P. J. & Eitan, S. Socially induced morphine pseudosensitization in adolescent mice. Behav Pharmacol 21, 112–120 (2010).

54. Ziółkowska, B., Gieryk, A., Solecki, W. & Przewłocki, R. et al. Temporal and anatomic patterns of immediate-early gene expression in the forebrain of C57BL/6 and DBA/2 mice after morphine administration. Neuroscience 284, 107–124 (2015).

55. McReynolds, J. R., Christianson, J. P., Blacktop, J. M. & Mantsch, J. R. What does the Fos say? Using Fos-based approaches to understand the contribution of stress to substance use disorders. Neurobiol Stress 9, 271–285 (2018).

56. Tyson, A. L. & Margrie, T. W. Mesoscale microscopy and image analysis tools for understanding the brain. Prog Biophys Mol Biol 168, 81–93 (2022).

57. Susaki, E. A. et al. Advanced CUBIC protocols for whole-brain and whole-body clearing and imaging. Nat Protoc 10, 1709–1727 (2015).

58. Ertürk, A. et al. Three-dimensional imaging of solvent-cleared organs using 3DISCO. Nat. Protocols 7, 1983–1995 (2012).

59. Murray, E. et al. Simple, Scalable Proteomic Imaging for High-Dimensional Profiling of Intact Systems. Cell 163, 1500–1514 (2015).

60. Lehtinen, J. et al. Noise2Noise: Learning Image Restoration without Clean Data. Preprint at 10.48550/ARXIV.1803.04189 (2018).

61. Tyson, A. L. et al. A deep learning algorithm for 3D cell detection in whole mouse brain image datasets. PLOS Computational Biology 17, e1009074 (2021).

62. Eichenberger, B. T., Zhan, Y., Rempfler, M., Giorgetti, L. & Chao, J. A. deepBlink: threshold-independent detection and localization of diffraction-limited spots. Nucleic Acids Res 49, 7292–7297 (2021).

63. Ronneberger, O., Fischer, P. & Brox, T. U-Net: Convolutional Networks for Biomedical Image Segmentation. Preprint at 10.48550/ARXIV.1505.04597 (2015).

64. Ester, M., Kriegel, H.-P., Sander, J. & Xu, X. A Density-Based Algorithm for Discovering Clusters in Large Spatial Databases with Noise. 226–331 (1996).

65. He, K., Zhang, X., Ren, S. & Sun, J. Deep Residual Learning for Image Recognition. Preprint at 10.48550/ARXIV.1512.03385 (2015).

66. Goutte, C. & Gaussier, E. A Probabilistic Interpretation of Precision, Recall and F-Score, with Implication for Evaluation. In Advances in Information Retrieval (eds. Losada, D. E. & Fernández-Luna, J. M.) vol. 3408 345–359 (Springer Berlin Heidelberg, Berlin, Heidelberg, 2005).

67. Howard, J. & Gugger, S. fastai: A Layered API for Deep Learning. (2020) doi:10.48550/ARXIV.2002.04688.

68. Haase, R. haesleinhuepf/apoc. (2025).

69. Breiman, L. Random Forests. Machine Learning 45, 5–32 (2001).

70. Tyson, A. L. et al. Accurate determination of marker location within whole-brain microscopy images. Sci Rep 12, 867 (2022).

71. Perez, F. & Granger, B. E. IPython: A System for Interactive Scientific Computing. Comput. Sci. Eng. 9, 21–29 (2007).

72. Claudi, F. et al. Visualizing anatomically registered data with brainrender. Elife 10, e65751 (2021).

73. Inc, P. T. Collaborative data science. https://plot.ly (2015).

74. Waskom, M. et al. mwaskom/seaborn: v0.8.1 (September 2017). Zenodo 10.5281/ZENODO.883859 (2017).

75. Sofroniew, N. et al. napari: a multi-dimensional image viewer for Python. Zenodo 10.5281/ZENODO.3555620 (2025).

76. Virtanen, P. et al. SciPy 1.0: Fundamental Algorithms for Scientific Computing in Python. Nature Methods 17, 261– 272 (2020).

